# CGGBP1-dependent CTCF-binding sites restrict ectopic transcription

**DOI:** 10.1101/2020.11.09.375782

**Authors:** Divyesh Patel, Manthan Patel, Subhamoy Datta, Umashankar Singh

## Abstract

Binding sites of the chromatin regulator protein CTCF function as important landmarks in the human genome. The recently characterized CTCF-binding sites at LINE-1 repeats depend on another repeat-regulatory protein CGGBP1. These CGGBP1-dependent CTCF-binding sites serve as potential barrier elements for epigenetic marks such as H3K9me3. Such CTCF-binding sites are associated with asymmetric H3K9me3 levels as well as RNA levels in their flanks. The functions of these CGGBP1-dependent CTCF-binding sites remain unknown. By performing targeted studies on candidate CGGBP1-dependent CTCF-binding sites cloned in an SV40 promoter-enhancer episomal system we show that these regions act as inhibitors of ectopic transcription from the SV40 promoter. CGGBP1-dependent CTCF-binding sites that recapitulate their genomic function of loss of CTCF binding upon CGGBP1 depletion and H3K9me3 asymmetry in immediate flanks are also the ones that show the strongest inhibition of ectopic transcription. By performing a series of strand-specific reverse transcription PCRs we demonstrate that this ectopic transcription results in the synthesis of RNA from the SV40 promoter in a direction opposite to the downstream reporter gene in a strand-specific manner. The unleashing of the bidirectionality of the SV40 promoter activity and a breach of the transcription barrier seems to depend on depletion of CGGBP1 and loss of CTCF binding proximal to the SV40 promoter. RNA-sequencing reveals that CGGBP1-regulated CTCF-binding sites act as barriers to transcription at multiple locations genomewide. These findings suggest a role of CGGBP1-dependent binding sites in restricting ectopic transcription.

## Introduction

CTCF binding sites serve as insulator and chromatin barrier elements (Phillips and Corces 2009; Ong and Corces 2014). CTCF binds to insulators and organizes the genome in chromatin regulatory domains. Regulatory elements within one chromatin regulatory domain interact more frequently than regulatory elements across the domain (Van Bortle et al. 2012; Nichols and Corces 2015; Arzate-Mejía et al. 2018). By partitioning the genome into regulatory domains, CTCF, along with cohesin, establishes cell-type-specific gene expression patterns (Ren and Zhao 2019; Hou et al. 2010). The two loci pioneering the characterisation of the insulator function of CTCF were the beta-globin locus control region and Igf2-H19 locus (Han et al. 2008; Ulaner et al. 2003; Valadez-Graham 2004; Farrell et al. 2002; Kurukuti et al. 2006). CTCF-binding sites, along with those of other associated chromatin regulatory proteins, serve as the boundaries of the chromatin loops and higher-order chromatin structures such as Topologically Associated Domains (TADs) (Narendra et al. 2016; Giorgetti et al. 2016; Dekker and Mirny 2016). CTCF thus allows the enhancer-promoter communication within TADs and prevents the inter-TAD interaction of the gene-regulatory elements (Dekker and Mirny 2016; Galupa and Crocker 2020; Ghirlando and Felsenfeld 2016). Further, CTCF acts as a barrier element by marking the boundaries of heterochromatin domains that are enriched with repressive histone marks such as constitutive silencing H3K9me3 (may bound by HP1) and temporary silencing marks H3K27me3 (may bound by polycomb repressors) (Lu et al. 2016; Kim et al. 2011; Barkess and West 2012). CTCF also prevents the spread of heterochromatin into the gene-rich euchromatin (Barkess and West 2012; Cuddapah et al. 2009).

A subset of CTCF-binding sites is dependent on a less well studied protein CGGBP1 (Patel et al. 2020, 2019). Recently described CGGBP1-dependent CTCF-binding sites function as chromatin barrier elements (Patel et al. 2019). They restrict H3K9me3 signal spread and function as boundaries of H3K9me3-rich and H3K9me3 depleted regions (Patel et al. 2019). This property of CGGBP1-dependent CTCF-binding sites seems to not affect H3K27me3 and H3K4me3 levels (Patel et al. 2019) as much. H3K9me3 is a major gene silencing epigenetic mark which is also employed for chromatin compaction and prevention of noisy transcription. This epigenetic mark is predominant in the Giemsa-positive gene-poor regions and hence aids in condensation of such DNA and rendering it unavailable for active transcription (Becker et al. 2016; Ninova et al. 2019). H3K9me3 plays a prominent role in repeat silencing and retrotransposon inactivation (van Kruijsbergen et al. 2017; Bulut-Karslioglu et al. 2014). The different kinds of repeats marked by H3K9me3 include satellite repeats, tandem repeats and DNA transposons (Bulut-Karslioglu et al. 2014; Iglesias and Moazed 2017). H3K9me3 plays a vital role in maintaining genomic integrity by preventing the genomic integration of the LTR and non-LTR retrotransposons specifically interspersed repeats such as Alu-SINEs and L1-LINEs (Bulut-Karslioglu et al. 2014; van Kruijsbergen et al. 2017). Interestingly, the CGGBP1-dependent CTCF-binding sites are also mostly L1 repeat-rich and CTCF motif-poor. CTCF occupancy at these sites depends on CGGBP1 levels (Patel et al. 2019). This regulation of CTCF binding to repeat-derived binding sites by CGGBP1 does not seem to require the formation of a protein-protein complex between the two proteins. A comparison of CTCF occupancy between three different CGGBP1 levels (normal, depleted and overexpressed) suggests that there is cooperative facilitation of CTCF-repeat binding by CGGBP1 (Patel et al. 2019). However, since CGGBP1 itself is a repeat binding protein, this cooperativity of CGGBP1-CTCF binding is lost upon CGGBP1 overexpression just the way it is lost upon CGGBP1 depletion (Patel et al. 2019). The repeat-origins of CTCF-binding sites are established. In the primates, Alu-SINEs have diverged into a large number of CTCF-binding sites in the human genome (Schmidt et al. 2012). However, CGGBP1-dependent CTCF-binding sites seem to have a preference for L1-LINEs. Remarkably, the CGGBP1-dependent CTCF-binding sites in L1 repeats concentrate on motif-like subsequences that are common between Alus and L1 elements (Patel et al. 2019). Thus, even with disparate evolutionary origins, L1 and Alu repeat function as sites where the CGGBP1-CTCF axis operates to regulate the patterns of H3K9me3 patterns. The functional significance of the CGGBP1-dependent CTCF-binding sites however remains unclear. This is especially interesting given the evolutionary unrelatedness of CTCF and CGGBP1. The former is conserved in vertebrates whereas the latter is present only in amniotes. One significance of the CGGBP1 regulation of CTCF binding is highlighted by our recent work that CGGBP1 levels regulate cytosine methylation at CTCF-binding motifs in a non-stochastic manner (Patel et al. 2018, 2020).

The proteins with which CTCF and CGGBP1 form complexes shed some light on the possible functions of CGGBP1-dependent CTCF-binding sites and the mechanisms through which they are regulated. CGGBP1 itself is a component of the enhancer-binding proteins complexes containing YY1 and CTCF (Weintraub et al. 2017). The histone methyltransferases SUV4 and SUV39 family member enzymes form complexes with cohesin ring family members and thus associate with CTCF (Hahn et al. 2013). HMT SUV39H2 is also associated with CGGBP1 (Singh et al. 2011; Singh and Westermark 2015). Although a fraction of CTCF and CGGBP1 do co-immunoprecipitate with each other, such indirect interactions seem to direct the coregulation between CTCF and CGGBP1 at CGGBP1-dependent CTCF-binding sites (Patel et al. 2019). Nucleophosmin forms complexes with both CTCF and CGGBP1 (Yusufzai et al. 2004; Hein et al. 2015). Interestingly, boundaries of the L1-rich lamina associated domains show contrasting levels of CTCF occupancy outside and within the LAD. This LAD boundary specific pattern also depends on the functions of CGGBP1. However, the CTCF-CGGBP1 complexes detected *in situ* do not localize to the nuclear periphery (Patel et al. 2019).

CGGBP1 has been implicated in the regulation of transcription of interspersed repeats. It is also required for normal RNA Polymerase 2 activity. Upon growth stimulation of normal human fibroblasts, the transcript elongation by RNA Polymerase 2 seems to depend on the levels of CGGBP1 (Patel et al. 2019; Agarwal et al. 2016; Ichiyanagi 2014; Cardiello et al. 2014; Singh and Westermark 2015). Circumstantial evidence suggests that this H3K9me3 asymmetry in the flanks of CGGBP1-dependent CTCF-binding sites may lead to asymmetrical RNA abundance in 10 kb long flanking regions (Patel et al. 2019). CTCF binding sites exert a transcription repressive effect in *cis*. CTCF was first reported as a transcriptional repressor of the chicken c-myc gene (Lobanenkov et al. 1990; Holwerda and de Laat 2013). Further studies have found transcriptional activator activity of the CTCF (Klenova et al. 1993; Lobanenkov et al. 1990). The regulatory function of CTCF binding sites is determined by its interacting proteins, the location of CTCF binding sites relative to the transcription site of a gene (Holwerda and de Laat 2013; Nishana et al. 2020; Lobanenkov et al. 1990).

Thus, there seems to be a functional link between CGGBP1-dependent CTCF-binding sites, regulation of specific histone modifications including H3K9me3 and RNA Polymerase 2 activity. Here we have investigated the effects CGGBP1 levels exerts on how the candidate CGGBP1-dependent CTCF-binding sites affect histone modifications, transcript levels and RNA Polymerase 2 occupancy. By using an episomal system that drives transcription from the SV40 promoter under the control of the SV40 enhancer, we show that CGGBP1 depletion promotes ectopic transcription activity from the SV40 promoter. We report that the early transcriptional activity downstream of the SV40 promoter remains unaffected by CGGBP1 depletion. However, the upstream late transcription, which normally depends on the SV40 enhancer, is induced upon CGGBP1 depletion. This generates an ectopic strand-specific transcription from the SV40 promoter. By using different CTCF-binding sites juxtaposed with the SV40 promoter we have found that this effect of CGGBP1 depletion is linked to the presence of CGGBP1-dependent CTCF-binding sites. These results shed light on the mechanisms of action of CGGBP1-dependent CTCF-binding sites and highlight the role of CGGBP1 in transcription regulation that can bypass the enhancer-dependence of promoters.

## Results and Discussion

### Genomic CGGBP1-dependent CTCF-binding sites and characterization of LoB5, a candidate CGGBP1-dependent CTCF-binding site

The locations of the 879 CGGBP1-dependent CTCF-binding sites were analyzed with respect to the known TSSs (Fantom database). These CGGBP1-dependent CTCF-binding sites were located at long distances from the TSSs (185 kb ± 388.021 kb) in the gene-poor regions and did not seem to be involved in *cis*-regulation of gene expression. We could detect transcripts generated from some of these TSS pairs of the same genes (such that the TSS pairs were located on either side of the CGGBP1-dependent CTCF-binding site) which were the closest to the CGGBP1-dependent CTCF-binding sites (Table 1). However, there was no consistent effect of CGGBP1 depletion on transcript levels of these genes (Figure 1A). A survey of transcript levels derived from five such TSSs revealed that although some pairs of TSSs do display a differential activity of the two TSSs in CT that is lost in KD, such an effect was not observed consistently in the selected TSS pairs (Figure 1A). For some genes (NRXN2 and OPRL) the transcript level differences were in line with the RNA Polymerase occupancy observed at the same regions suggesting that to some extent the transcript levels reflect the activity of RNA Polymerase 2 at these TSSs (Figure 1B).

**Table 1:** The selected alternative TSS pairs of genes with the least distances to the CG-GBP1-regulated CTCF-binding sites. The table shows the gene symbols and the locations of the TSSs from the nearest CGGBP1-regulated CTCF-binding sites. Table represents the distance between CGGBP1 regulated chromatin barrier element and permissive TSSs used for PCR assay.

| Genes with TSSs separated by CGGBP1-dependent CTCF-binding site | Type of change in CTCF-binding upon CGGBP1 depletion | Upstream distance between the TSS and CTCF-binding site (kb) | Downstream distance between the TSS and CTCF-binding site (kb) |
| --- | --- | --- | --- |
| NRXN2 | Loss | 21.5 | 13.6 |
| PBX1 | Loss | 13.5 | 8.1 |
| OPRL | Loss | 1.6 | 2.8 |
| SPTBN1 | Gain | 12.7 | 87.7 |
| COMT1 | Gain | 5.8 | 2.9 |

**Figure 1:**
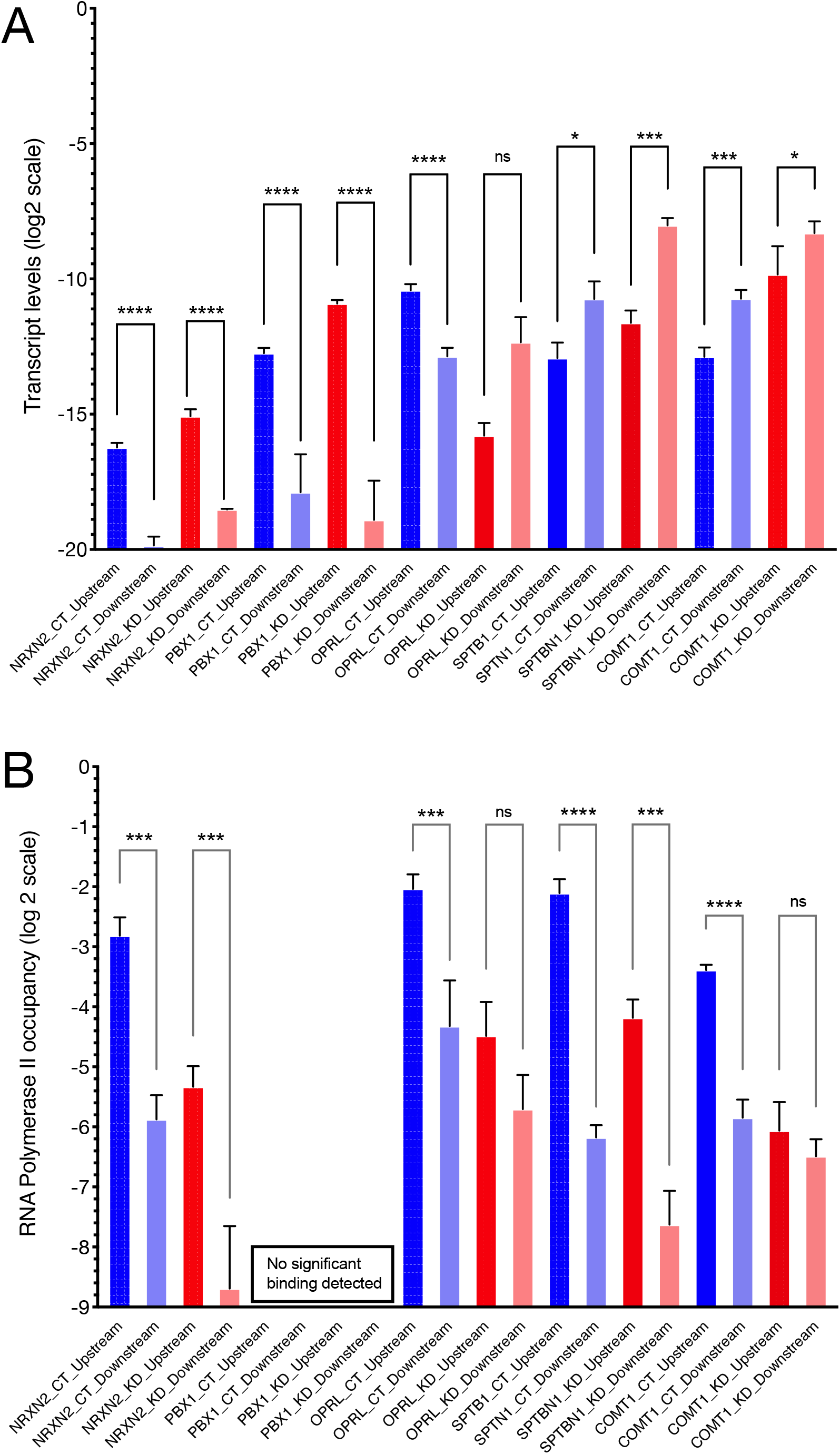
RNA levels and RNA Polymerase 2 occupancy at the TSSs of selected genes tabulated in Table 1. **A:** The RNA levels at the TSSs of the genes are calculated by the double delta Ct method. Beta-actin (ACTB) has been used as a quantitation control in all the PCRs. The Y-axis shows the ddCt values on a log2 scale. The location of the TSSs relative to the nearest CGGBP1-regulated CTCF-binding sites are indicated by suffixes “upstream” or “downstream”. **B:** RNA Polymerase 2 occupancy at the same TSSs as shown in **A** shows that apart from NRXN2, none of the TSSs showed a correspondence between RNA levels and RNA Polymerase 2 occupancy. For PBX1 the Ct values obtained were too low to be used reliably for ddCt analysis and were eliminated. The levels of amplification from the input DNA was used as a quantitation control against respective ChIP. The Y-axis shows the ddCT values on a log2 scale.

One of the known functions of these CTCF-binding sites is the maintenance of differential levels of H3K9me3 in the flanks such that the differences depend on the levels of CGGBP1. The *cis* length range in which H3K9me3 asymmetry has been studied earlier is 10 kb. It was not clear however if these TSSs were under the influence of H3K9me3 silencing or not. We selected some CGGBP1-dependent CTCF-binding sites with the highest reported asymmetries in the cumulative H3K9me3 signals in their 10 kb flanks and verified two important parameters at them for further studies: (i) CGGBP1-dependence of CTCF occupancy at them, and (ii) H3K9me3 asymmetry in their immediate flanks. Three of these regions which we pursued are called LoB3, LoB5 and GoB4. The genomic locations and contexts of these CGGBP1-dependent CTCF-binding sites are shown in figure S1 (Fig S1, A to C). We identified one region, called LoB5, where we could consistently perform specific PCR amplification. At LoB5, CTCF occupancy was lost upon CGGBP1 depletion (Fig 2A). The other two regions LoB3 and GoB4 showed a weaker loss and gain of CTCF binding respectively upon CGGBP1 depletion (Fig 2, B and C). Also, the H3K9me3 levels were asymmetric in LoB5 flanks only in the presence of CGGBP1 and were lost upon CGGBP1 depletion (Fig 2D). The immediate flank transcription at LoB5 also exhibited expected asymmetry in CT as well as KD (not shown). Although GoB4 and LoB3 showed some CGGBP1-dependence of CTCF binding, they did not recapitulate the expected H3K9me3 levels in flanks (Fig 2, E and F). Thus, LoB5 presented us with a region where the CTCF and H3K9me3 ChIP-seq data could be independently verified. These findings suggested that LoB5 could be used as a model region to explore the role of the CTCF-CGGBP1 axis at such candidate barrier elements. We cloned LoB5 in an episomal vector system pGL3-Control. The LoB5 element was inserted upstream of the SV40 promoter in *KpnI-XhoI* sites (Fig 3A; the figure also shows the locations of the various regions and primer binding sites used further on in various experiments). Unlike the endogenous LoB5 locus that is distant from TSSs, in this construct, the LoB5 element was artificially juxtaposed against an RNA Polymerase 2 promoter. We tested if the LoB5 element in the episomal system retained its properties of CGGBP1-dependent CTCF binding and H3K9me3 asymmetry in immediate flanks. ChIP-qPCRs revealed that LoB5 is a potent CTCF-binding site in CT. Upon CGGBP1 knockdown CTCF-binding at episomal LoB5 was lost (Fig 3B). This was verified using primer pairs that amplified the endogenous LoB5 and episomal LoB5 exclusively or commonly. Similarly, constructs containing LoB3 and GoB4 were also subjected to CTCF ChIP-qPCRs in CT and KD but the expected loss of binding of CTCF upon CGGBP1 depletion was not observed on those clones (Fig 3, C and D). The H3K9me3 levels in the upstream and downstream regions of episomal LoB5 showed the same effect of CGGBP1 depletion as was observed for the genomic endogenous LoB5 locus (Fig 3E). Again for LoB3 and GoB4 constructs, the H3K9me3 levels in the flanks of the inserts did not mimic the expected genomic H3K9me3 patterns (Fig 3, F and G). We thus focussed on the LoB5 construct as an episomal system to study the functions of LoB5 as a model CGGBP1-dependent CTCF-binding site.

**Figure 2:**
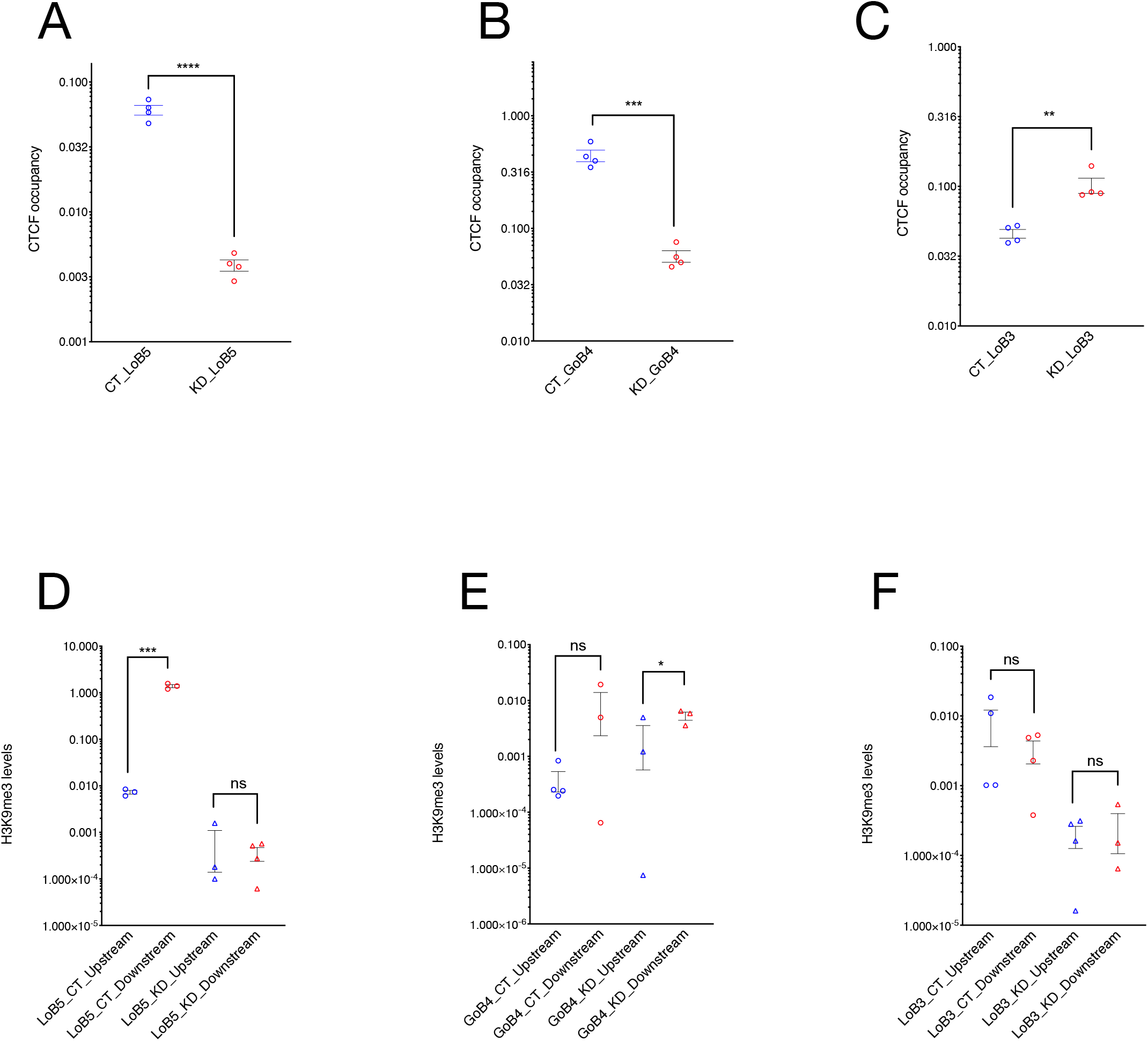
LoB5 shows the expected loss of CTCF occupancy and H3K9me3 asymmetry upon CGGBP1 depletion. **A-C:** LoB5, GoB4 and LoB3 sites were cloned in pGL3-control vector. The cloned plasmid was transfected in HEK293T cells with normal and depleted levels of CGGBP1 separately. Total CTCF occupancy at CTCF binding sites (Endogenous and episomal) was compared between CT and KD for LoB5, GoB4 and LOB3. The Ct value for CTCF enrichment was normalised with input and CTCF enrichment are plotted in arbitrary units. Statistical significance was determined by using the unpaired t-test (p-value = 0.05). CTCF binding is strongly reduced at the LoB5 site in KD **(A)**. Depletion of CGGBP1 leads to decreased occupancy of CTCF at the GoB4 site **(B)** and increased occupancy at the LoB3 site **(C)**. **D-F:** H3K9me3 levels in the immediate flanks of the endogenous CTCF binding sites were compared between CT and KD. The Ct value for H3K9me3 enrichment was normalised with input and H3K9me3 enrichment is represented in arbitrary units (Y-axis). Statistical significance was determined by using the unpaired t-test (p-value = 0.05). H3K9me3 levels show significant asymmetry in the immediate flanks of the endogenous LoB5 CTCF binding site in CT (p-value < 0.05). The decrease in H3K9me3 levels in the downstream region causes loss of H3K9me3 asymmetry in KD **(D)**. A non-significant asymmetry in H3K9me3 levels is observed in the immediate downstream flanks of the GoB4 CTCF binding site in CT. However, H3K9me3 levels show mild but significant asymmetry in KD **(E)**. The immediate flanks of the LoB3 sites maintained similar levels of H3K9me3 in CT and KD **(F)**.

**Figure 3:**
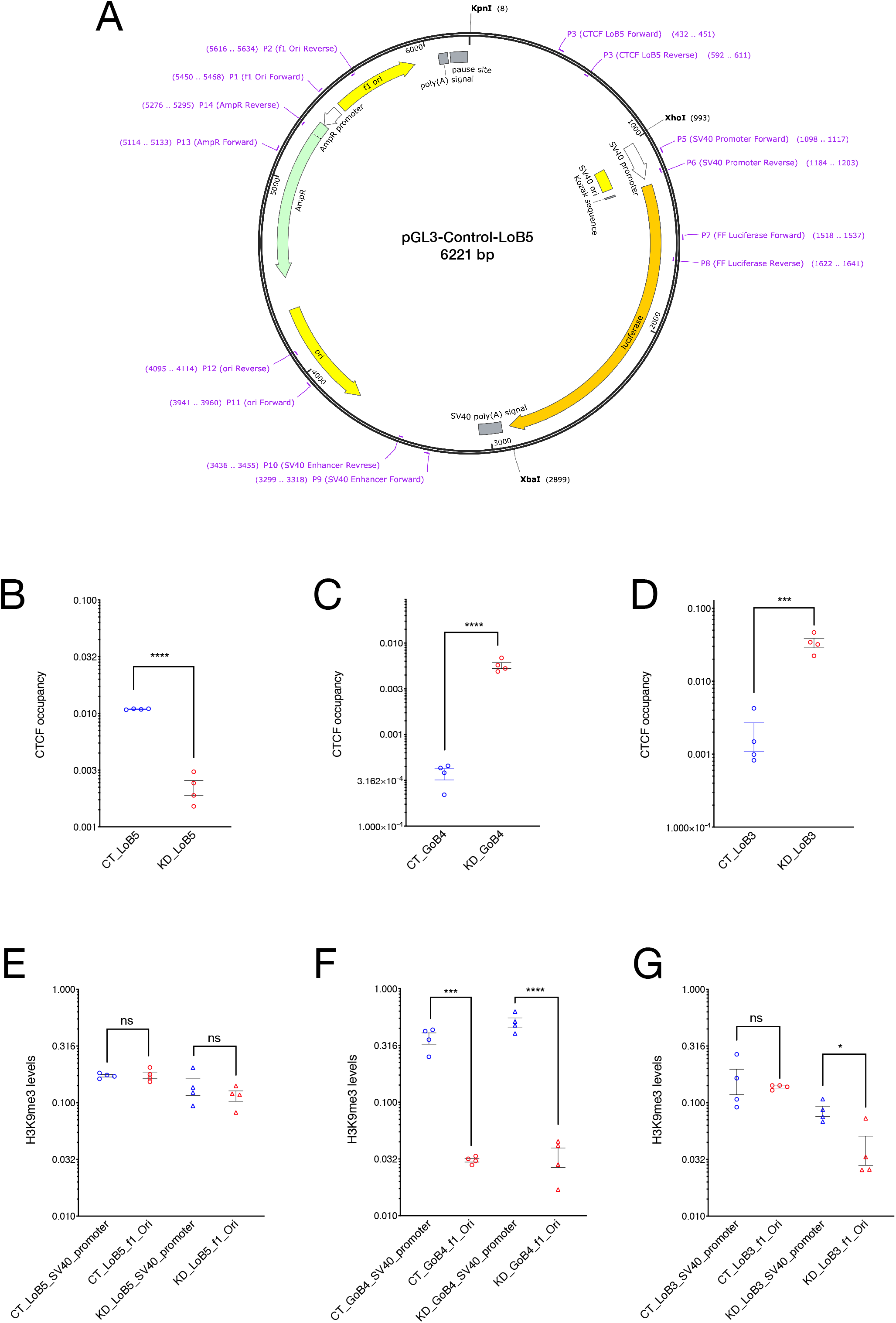
Characterisation of CGGBP1-regulated CTCF binding sites in the episomal system. **A:** Schematic representation of the pGL3-control episomal vector system. Candidate LoB and GoB sites along with approximately 250 bp flanking sequences were cloned upstream of the SV40 promoter. The schematic shows the LoB5 CTCF binding site along with primers used to determine CTCF binding, transcript levels, RNA-Polymerase II, H3K9me3 and H3K27me3 levels at different regions of the episome. **B-D:** CTCF occupancy at episomal LoB and GoB sites were compared between CT and KD by using primer P3 (cloned CTCF sites forward) and P6 (SV40 promoter reverse). The Ct value for CTCF enrichment was normalised with input and CTCF enrichment are plotted in arbitrary units. Statistical significance was determined by using the unpaired t-test (p-value = 0.05). Episomal LoB5 CTCF binding has shown a strong decrease in CTCF binding in KD **(B)**. CTCF occupancy at the episomal GoB4 site increases significantly **(C)**, similarly, the LoB3 site in the episome shows a significant increase in CTCF binding upon CGGBP1 depletion **(D)**. **E-G:** H3K9me3 levels at f1Ori and SV40 promoter located upstream and downstream respectively of the episomal CTCF binding sites were compared between CT and KD. The Ct value for H3K9me3 enrichment was normalised with input and H3K9me3 levels are represented as arbitrary units (y-axis). Statistical significance was determined by using the unpaired t-test (p-value = 0.05). H3K9me3 levels are comparable upstream and downstream of the episomal LoB5 site in CT and KD **(E)**. The upstream (f1Ori) of the episomal GoB4 CTCF binding site display comparatively reduced levels of the H3K9me3 than the SV40 promoter in CT. The asymmetric distribution of the H3K9me3 is potentiated in KD **(F)**. Comparable levels of H3K9me3 are observed at f1Ori and SV40 promoter in CT. A strong decrease in H3K9me3 levels at f1Ori increases asymmetry of H3K9me3 levels in immediate flanks of episomal LoB3 site in KD.

### Regulation of LoB5-SV40 promoter activity by CGGBP1

We next tested if CGGBP1 levels affected the LoB5-SV40 promoter activity. In this episomal system, the SV40 promoter activity is driven by an upstream enhancer located ~2 kb away (Fig 3A) (Kadesch and Berg 1986; Benoist and Chambon 1981; Shaw et al. 1985; Sassone-Corsi et al. 1984; Kelly and Wildeman 1991). Compared to the empty vector (SV40 promoter), the LoB5-SV40 promoter did not show any significant difference in the promoter activity as measured by Firefly Luciferase activity (Fig S2). In these experiments, we used Renilla Luciferase as an internal control for the normalization of systemic variables (Fig S2).

The SV40 promoter is a bidirectional promoter (Byrne et al. 1983; Hertz and Mertz 1988). Its basal downstream activity is driven by the 21 bp element that contains the CAAAT box and is proximal to the TSS (Byrne et al. 1983; Hertz and Mertz 1988). The upstream transcriptional activity is dependent on the 70 bp repeat elements located distal to the TSS (Wasylyk et al. 1983). We first measured the transcript levels of the Luciferase gene. Luciferase transcript levels were not changed by CGGBP1 depletion (Fig S3A). Since CGGBP1 depletion also caused a loss of CTCF binding, these findings reinforced that a CGGBP1, and potentially CTCF as well, do not act as *cis* regulators of SV40 promoter activity. The upstream enhancer drives the SV40 promoter (Hertz and Mertz 1988) and insertion of foreign DNA upstream of the SV40 promoter in pGL3-Control has been used as a tool to identify potential insulator sequences which can block the communication between enhancer and the promoter. However, these findings also suggested that there is no insulator-like activity of the LoB5 element in the episomal system.

Next, we measured transcript levels from various other regions of the episome. We found that just like the Luciferase gene, at the LoB5-SV40 promoter, AmpR gene, Ori and f1Ori there were no significant changes in transcript levels in KD compared to CT (Fig S3B). However, the SV40 enhancer showed a mild decrease (Fig S3B).

We further investigated if CGGBP1 depletion affected the occupancy of RNA Polymerase 2 in all these regions. Unexpectedly, the RNA Polymerase 2 occupancy was reduced in KD as compared to CT at all the regions except the LoB5-SV40 promoter and the Luciferase gene (Fig S3C). We then calculated a ratio of transcript abundance and RNA Polymerase 2 occupancy to gauge the transcript productivity of RNA Polymerase 2 presence on the DNA. We found that f1Ori had the highest transcript abundance to RNA Polymerase 2 occupancy ratio followed by the SV40 enhancer (derived from data shown in Fig S3, B and C; not shown). This indicated that upon CGGBP1 depletion, transcription activity at f1Ori increases. At the same time, the coupling between the SV40 enhancer and the LoB5-SV40 promoter, which is expected to drive the promoter activity, was not retained in KD. Upon CGGBP1 knockdown, the increase in the transcriptionally active RNA Polymerase 2 presence at SV40 enhancer did not produce a concomitant increase in Luciferase gene transcript levels. However, a highly similar increase in RNA Polymerase 2 occupancy and transcript abundance at f1Ori and SV40 Enhancer suggested that upon CGGBP1 depletion, the SV40 Enhancer and LoB5-SV40 promoter couple to drive transcription upstream towards the f1Ori.

### Restriction of bidirectional transcription from LoB5-SV40 promoter by CGGBP1

For the possibility of transcription of f1Ori by LoB5-SV40 promoter to materialize, the RNA Polymerase 2 activity from SV40 promoter must overcome the transcription pause signal located between the MCS and the f1Ori (Fig 3A). Thereby, transcripts must be generated that span the region from the LoB5-SV40 promoter to the f1Ori.

PCRs on randomly primed cDNA using primers located in f1Ori and SV40 revealed that transcripts spanning f1Ori and SV40 promoter are formed for LoB3, LoB5 as well as GoB4 (Fig S4A). The amplification was observed from DNase-digested RNA templates but not observed upon digestion with RNaseH or RNaseI (not shown). Interestingly, the CGGBP1-dependence of this SV40 promoter-f1Ori transcript was observed very strongly for LoB5, weakly for GoB4 and not at all for LoB3 (Fig S4A). The same length of SV40 promoter-f1Ori transcript was obtained using oligo-dT for cDNA synthesis followed by PCR using primers located in f1Ori and SV40 Promoter (Fig S4B). These results suggested that the SV40 promoter-f1Ori transcript is polyadenylated at a location 3’ to the f1Ori. The depletion of CGGBP1 thus allows RNA Polymerase 2 to breach the transcription pause and polyA sites located between SV40 promoter and f1Ori (Fig s4A). As a control, we tested if the polyA signal and RNA Polymerase 2 pause site between the 3’ end of the Luciferase gene and SV40 enhancer is also breached. Using strand-specific cDNA synthesized using P5 or P10 (Fig S4A), we were not able to amplify any products using one primer in SV40 enhancer and another in SV40 promoter (not shown). Thus, the directionality of the SV40 promoter towards the Luciferase gene was not altered but ectopic transcription from the SV40 promoter through LoB5 towards f1Ori occurred upon CGGBP1 depletion.

To further characterize the direction of transcription of the SV40-LoB5-f1Ori we performed strand-specific PCRs. cDNA was prepared from the terminal primers located in f1Ori or SV40 promoter and the PCRs were performed using both these primers. The entire ~ 2 kb long product was amplified from cDNA synthesized using f1Ori forward primer only (Fig S4B). The cDNA generated using the SV40 promoter reverse primer did not give rise to any product (Fig S4B). The f1Ori forward primed cDNA showed stronger amplification of the SV40-LoB5-f1Ori product in KD and only minimal amplification was detected in CT (Fig S4B). The 1.97 kb long PCR product was sequence-verified using two opposite outgoing primers in the LoB5 (data not shown). Further, by using the cDNA generated using the f1Ori terminal primer, we performed PCRs for terminal fragments in the 2kb SV40-LoB5-f1Ori transcript (Fig S4C). The f1Ori levels were much less compared to the levels at the SV40 promoter (Fig S4C). These findings indicated that the SV40 promoter-f1Ori transcript synthesis begins at the SV40 promoter and due to incomplete synthesis of transcripts and truncations before f1Ori, the levels of the 3’ end of the transcript are lower than those at the 5 ‘end.

Together, these findings showed that upon CGGBP1 depletion the SV40 promoter activity is driven to synthesize the SV40-LoB5-f1Ori transcript that is in the opposite direction and on the strand complementary to the Luciferase gene sense strand. Interestingly, depletion of CGGBP1 allowed this atypical transcription to occur despite the presence of the TTS that normally restricts transcription. However, this ability of RNA Polymerase 2 to breach the pause site was observed only for the SV40-upstream region that contained LoB5. For the downstream TTS located between the Luciferase gene and SV40 enhancer, no ectopic transcription was observed.

The LoB5 element recapitulates some of its key endogenous properties in the episomal system. The LoB5 belongs to a set of regions that exhibit CGGBP1-dependent CTCF binding and are rich in L1 repeats. These findings raised the possibility that the SV40 T-antigen binding sites in the L1-rich CGGBP1-dependent CTCF-binding sites genome-wide could exhibit ectopic transcription in the absence of CGGBP1. L1 repeats and L1 repeat-derived sequences also constitute the primary CGGBP1-dependent CTCF-binding sites. To assess the extent to which CGGBP1-dependent CTCF-binding sites affect transcription boundaries at endogenous loci genome-wide, we performed RNA-seq on CT and KD. Our approach focussed on identifying transcription boundaries in CT that are breached in KD bidirectionally.

A comparison of CT and KD RNA profiles showed two prominent features. First, the weak short range transcription sites in CT (812 filtered sites) were transcribed into longer contiguous transcripts in KD albeit at a lower level showing a loss of transcription boundaries in KD (Fig 4, A and B). As a converse, at a filtered set of 403 sites with string transcription in CT, a restriction of transcription was observed in KD (Fig 4, C and D). Second, by comparing the transcription boundaries in CT that are breached in KD with CTCF occupancy in CT and KD, we observed that the transcription boundaries are restricted by specific CTCF occupancy patterns in CT. These CTCF binding patterns were disrupted in KD (Fig 4, E and F). The transcription start and end sites both exhibited a presence of CTCF occupancy immediately upstream of the point of transcription restriction in CT as compared to KD. These results suggested that transcription at ectopic transcription sites at multiple locations in the genome is restricted by CGGBP1-dependent CTCF-binding sites. These CGGBP1-dependent CTCF-binding sites are mainly the L1 repeats. Interestingly, the L1 repeats are also strong binding sites for the SV40-Large T antigen and transcription factors, such as SP1, which are required for bidirectional transcription (Gidoni et al. 1985; Gruss et al. 1988). Our results suggest that the regulation of bidirectional transcription from L1 repeats by SP1 reported earlier is a broader process regulated in part by the CGGBP1-CTCF axis as well.

**Figure 4:**
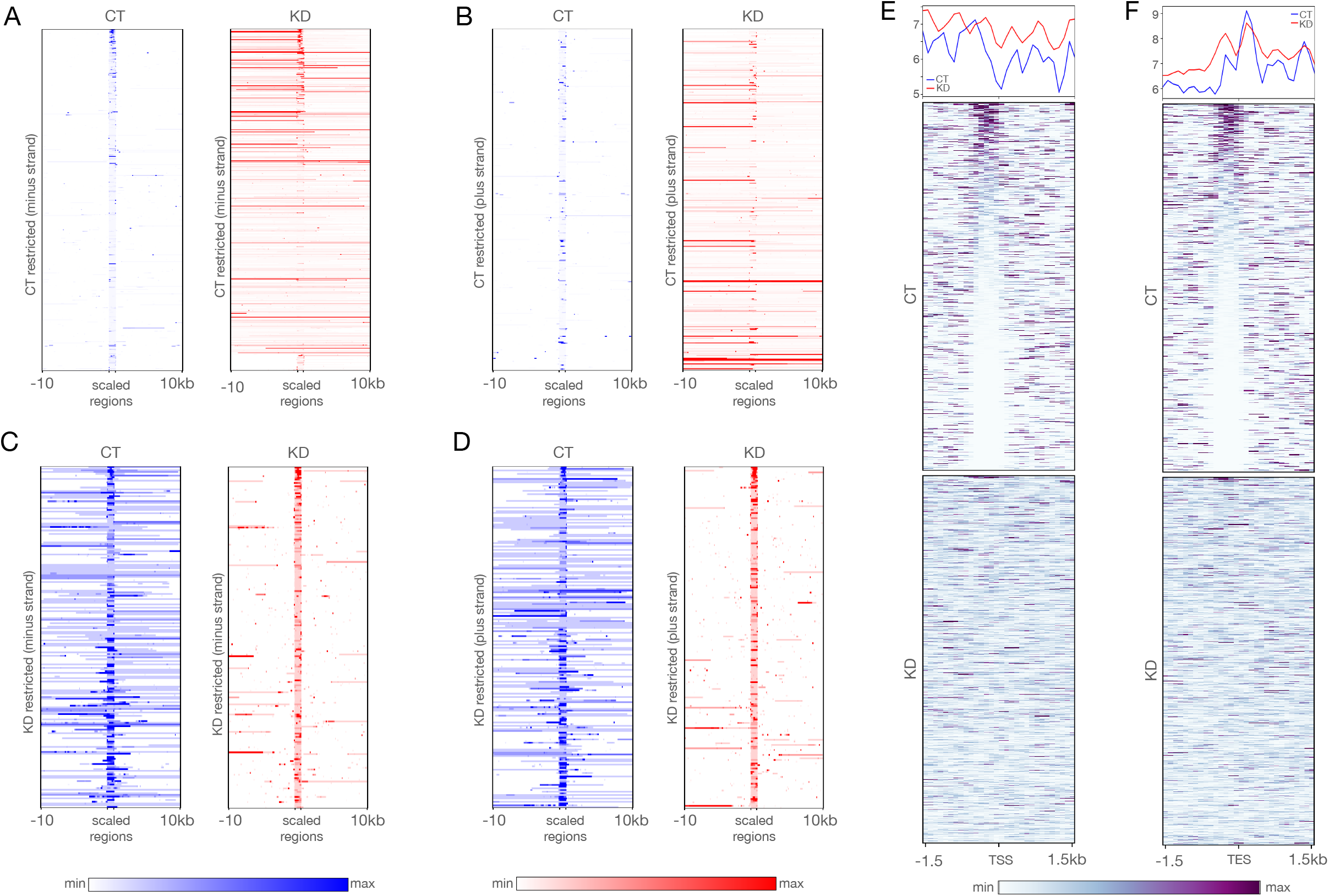
RNA-seq reveals CGGBP1-regulated CTCF binding patterns at transcription restriction sites. **(A** and **B)** The short-range weakly transcribing regions in CT (blue) showed stronger transcription in a longer range upon depletion of CGGBP1 in (KD, red) for both minus **(A)** and plus **(B)** strands respectively. **(C** and **D)** The regions of our genome which exhibit stronger and long-range transcription under normal levels of CGGBP1 (CT, blue) were found to be restricted to short-range transcription upon CGGBP1 knockdown (KD, red) on the minus **(C)** and plus **(D)** strands respectively. **(E** and **F)** The weakly transcribing regions of the genome in presence of CT were marked by the presence of CTCF-binding that acted as barrier to transcription in the upstream of the start **(E)** and end **(F)** sites of the transcripts which was not maintained in KD. The plots show a general disruption of CTCF binding pattern at these transcription restriction sites.

## Materials and Methods

### Cell culture

All the experiments on cells were performed in the HEK293T cells. These cells were cultured in serum (10% FBS) supplemented DMEM (AL007A). CGGBP1 depletion in these cells was achieved by lentiviral transduction of the lentiviral shmiR constructs (four different sites in the ORF) targeting CGGBP1 (KD) or Control-shmiR (CT) obtained from Origene as described earlier (PMID: 31547883) and not shown. The process of lentiviral production involved cotransfection of the third generation packaging plasmids (from Addgene): pRSV-Rev (12253), pMDLg/pRRE (12251) and pMD2.G (12259) and lentiviral constructs (from Origene) in equimolar ratios. For efficient transfection, Fugene (Promega) was used (3 μl/μg of DNA). Higher transduction yield was obtained with the help of Polybrene (Sigma), used at 1:10000 dilution of 10 mg/ml stock. Further, these cells were selected using 0.4 mg/ml of puromycin for stable transduction.

For the downstream experiments in HEK293T cells containing normal levels of CGGBP1 (CT) or depleted levels of CGGBP1 with the three different episomal constructs LoB5, LoB3 and GoB4 respectively along with the pRV-CMV plasmid (transfection control).

### Cloning of CGGBP1-dependent CTCF-binding sites into pGL3-Control vector

LoB5, LoB3 and GoB4 regions along with approximately 250 base flanking sequences were cloned between *KpnI* and *XhoI* restriction enzyme sites in the pGL3-control vector (PMID: 31547883). The LoB and GoB peaks and immediate flanking 250 base regions were amplified from genomic DNA of HEK293T cells by using primers described in Table 1. Clones were confirmed by Sanger sequencing (not shown). The PCR amplified genomic regions were double digested with *Kpn*I and *Xho*I. Similarly, the pGL3-Control vector was double digested with *Kpn*I and *Xho*I and cloned in pGL3-control. The primer sequences are mentioned below (5’-3’)

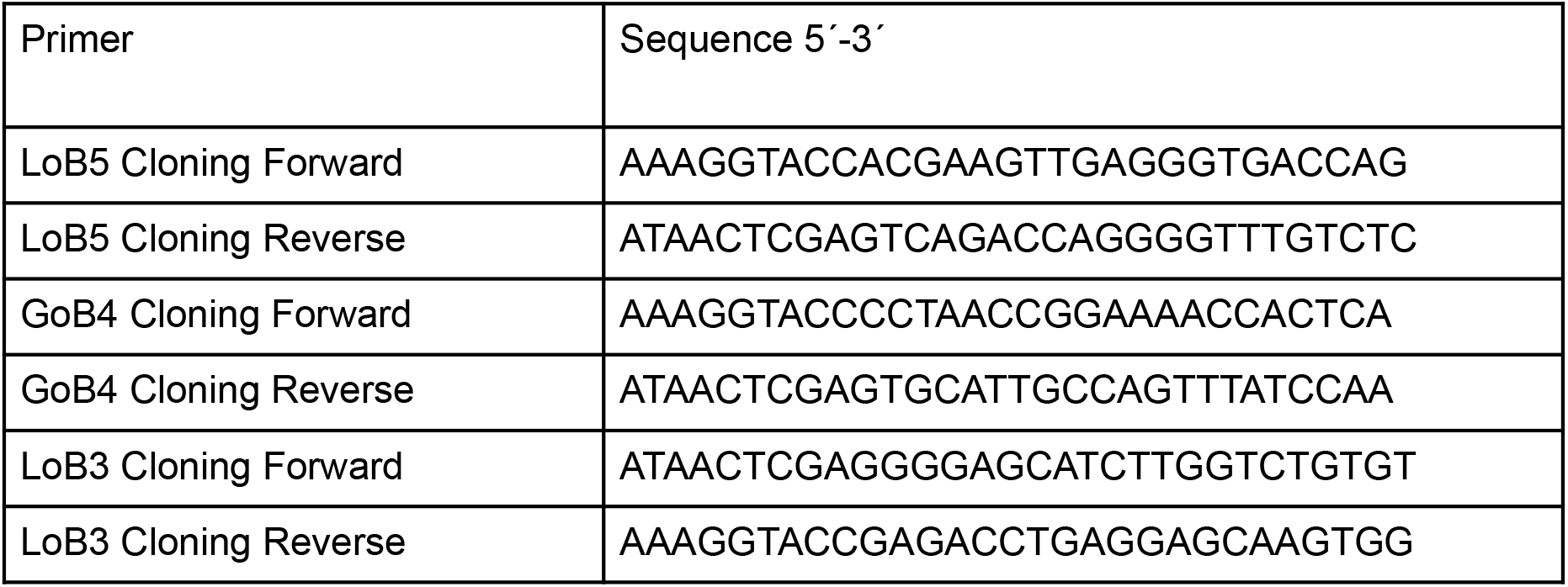

### Chromatin immunoprecipitation (ChIP)

Non-targeting or CGGBP1-targeting shRNA expressing lentivirus-transduced HEK293T cells (as described above) were grown in DMEM supplemented with 10% FBS. pGL3-LoB3, pGL3-LoB5 or pGL3-GoB4 constructs were transfected [3 μg per 10 cm plates] into CT and KD cells. A pRL-CMV plasmid (E2231, Promega) [2 μg per 10 cm plates] was co-transfected with these constructs as a transfection control. HEK293T cells were harvested after 48 h of transfection and subjected to RNA-isolation and ChIP-qPCR.

ChIP for the respective antibodies was performed as previously described (PMID: 31547883). Cells were harvested after crosslinking for 10 min at 37°C with 1% formaldehyde and quenching with Glycine (125 mM). Cells were washed with PBS twice and lysed in an SDS lysis buffer containing a cocktail of 1x protease phosphatase inhibitors (PI78441, Invitrogen). Based on our previous experience, the chromatin thus obtained was sonicated for 21 cycles of 30 seconds ON/30 seconds OFF to obtain DNA fragments in the range of 0.5 kb - 1 kb length. The clear fraction of sonicated chromatin was obtained after centrifugation (16000 rcf, 5 minutes, 4°C). 30 μl of fragmented chromatin was kept aside as input and the remaining (150 μl) was further diluted in ChIP-dilution buffer (0.01% SDS, 1.1% Triton X-100, 1.2 mM EDTA, 16.7 mM Tris-HCl, pH 8.1, 167 mM NaCl) containing 1X protease phosphatase inhibitors cocktail. The chromatin was pre-cleared by incubating with protein G sepharose beads for 4 hrs at 4°C followed by overnight incubation with a targeted primary antibody with mild tumbling at 4°C. Subsequently, the protein G sepharose beads were added and incubated for 1 hour followed by gentle centrifugation. The pelleted beads were washed with different buffers in the following order: low-salt IP wash buffer (0.1% SDS, 1% Triton X-100, 2 mM EDTA, 20 mM Tris–HCl and 150 mM NaCl), high-salt IP wash buffer (0.1% SDS, 1% Triton X-100, 2 mM EDTA, 20 mM Tris–HCl and 500 mM NaCl), LiCl IP wash buffer (0.25 M LiCl, 1% IGEPAL, 1% sodium deoxycholate, 1 mM EDTA and 10 mM Tris–HCl) and two washes of TE buffer (10 mM Tris–HCl and 1 mM EDTA). The immunoprecipitated DNA was eluted using an elution buffer (1% SDS and 0.1 M NaHCO3) and was reverse-crosslinked (addition of 20 μl of 5 M NaCl and incubation at 65°C for 6 hrs). Further, the DNA was subjected to Proteinase K (P2308, Sigma) digestion for 1 hour (addition of 10 μl of 0.5 M EDTA pH 8.0 and 20 μl of 1 M Tris-HCl pH 6.8 followed by 2 μl of 10 mg/ml Proteinase K. The DNA was purified following the column-based purification protocol (A1460, Promega). The set of different primary antibodies used in the ChIP assays are mentioned below:

| ChIP-target | Number of 10 cm plates<br>(about 20 million cells per plate) | Antibody used |
| --- | --- | --- |
| CTCF | 2 | 2.5 µg each of SC-28198 and SC-271514 SantaCruz) |
| RNA Polymerase 2 | 2 | 3 µg (05-623 Merck) |
| H3K9me3 | 1 | 3 µg (SC-130356 SantaCruz) |
| H3K27me3 | 1 | 2 µg of (ab6002 Abcam) |

### RNA extraction, cDNA synthesis and qRT-PCRs

RNA was isolated by using the TRIzol reagent as per the manufacturer’s instructions. RNA was isolated from pGL3-control construct transfected cells (as described above). Cells were washed with DEPC treated ice-chilled PBS twice and lysed by using TRIzol. Cells were scraped and cell lysate was collected in 1.5 ml Eppendorf. Chloroform (250 μl) was added to lysed cells and vigorously mixed and centrifuged at 10,000 rpm for 5 min. The aqueous layer was carefully transferred to the new tube and 550 μl isopropanol was added to the aqueous phase. The immunoprecipitated RNA was washed with 1 ml 75% ethanol in DEPC treated H2O and RNA was dissolved in 40 μl of DEPC treated water. The isolated RNA was digested with DNaseI (M0303S, 4 Units) at room temperature for 15 minutes. DNaseI digested RNA was isolated by using the TRIzol method.

### Luciferase assays

Lentiviral transduced HEK293T cells with normal and depleted levels of CGGBP1 were seeded in 96 well plates. LoB (pGL3-LoB3 and pGL3-LoB5), GoB (pGL3-GoB4) or pGL3-control empty vectors were transfected [100 ng per well] in CT and KD cells. The pRL-CMV plasmid (E2231, Promega) [25 ng per well] was co-transfected as transfection control. HEK293T cells were harvested after 72 h of transfection. The dual-luciferase assay was performed as per manufacturer protocol (E2920, Promega). Cells were washed with PBS and 100 μl of 1x Passive lysis buffer. 100 μl of Luciferase Assay Substrate resuspended in Luciferase Assay Buffer II was added to each well. Firefly luciferase activity was measured at 550 to 570 nm wavelength and followed by 100 μl of 1X Stop & Glo Reagent (part of the kit E1910, Promega) was added to each well. Renilla Luciferase activity was measured at 470 to 490 nm wavelength.

### cDNA synthesis

For cDNA synthesis using random primer the following method was used: cDNA synthesis was performed by using the SuperScript VILO cDNA Synthesis Kit (Invitrogen 11754050) as per the manufacturer’s protocol. RNA (2 μg) was incubated with 1X VILO Reaction Mix and 1X SuperScript Enzyme Mix. The reaction mix was incubated at room temperature for 15 minutes and followed by incubation at 42^0^ C for 60 minutes.

For cDNA synthesis using oligo-dT or primers P1 or P6 the following method was used: DNaseI digested RNA from LoB and GoB construct transfected cells CT and KD cells was used for cDNA synthesis. RNA ((2 μg) was incubated with primer at 65^0^ C for 5 minutes and snap-chilled immediately. M-MLV Reverse-transcriptase (Promega M1701, 200 Unit) in 1x reverse transcriptase buffer was added to primer annealed RNA and incubated at 40^0^ C for 60 minutes.

### Strand-specific PCR

cDNA synthesis by using an oligo-dT primer or strand-specific primer P1 (f1Ori Forward) and P6 (SV40 Promoter Reverse) were used as the template for PCR using primer P1 (f1Ori Forward) and P6 (SV40 Promoter Reverse). The specific amplifications were compared by Agarose gel electrophoresis.

### Sanger sequencing of PCR products

The PCR products obtained from the strand-specific PCR were subjected to the Sanger sequencing using sequencing primers P4 and P5 (Fig 4A).

### Statistical analysis

All the statistical analyses were performed on Graphpad Prism 8 and open office.

### Genome browser views

The genome-browser views of the three different CGGBP1-dependent CTCF-binding sites were obtained using Integrated Genomic Viewer (IGV). Repeat sequences were identified in 10kb flanks of the CGGBP1-dependent CTCF-binding sites by using RepeatMasker. The genomic coordinates of the identified repeat elements were used to generate bigwig signal files by using the deepTools tool.

### ChIP-qPCR

All quantitative PCR reactions were performed at 57°C annealing temperature and the specific template amplification was confirmed by agarose gel electrophoresis. Following are the PCR conditions: 95° C-5 minutes, (95° C-20 seconds, 55° C-20 seconds, 72° C-30 seconds, 80° C-30 seconds (signal capturing)) x45 followed by melting curve analysis (55 to 95 °C constant signal recording).

Following is the list of primers used for ChIP-qPCRs or RT-PCRs:

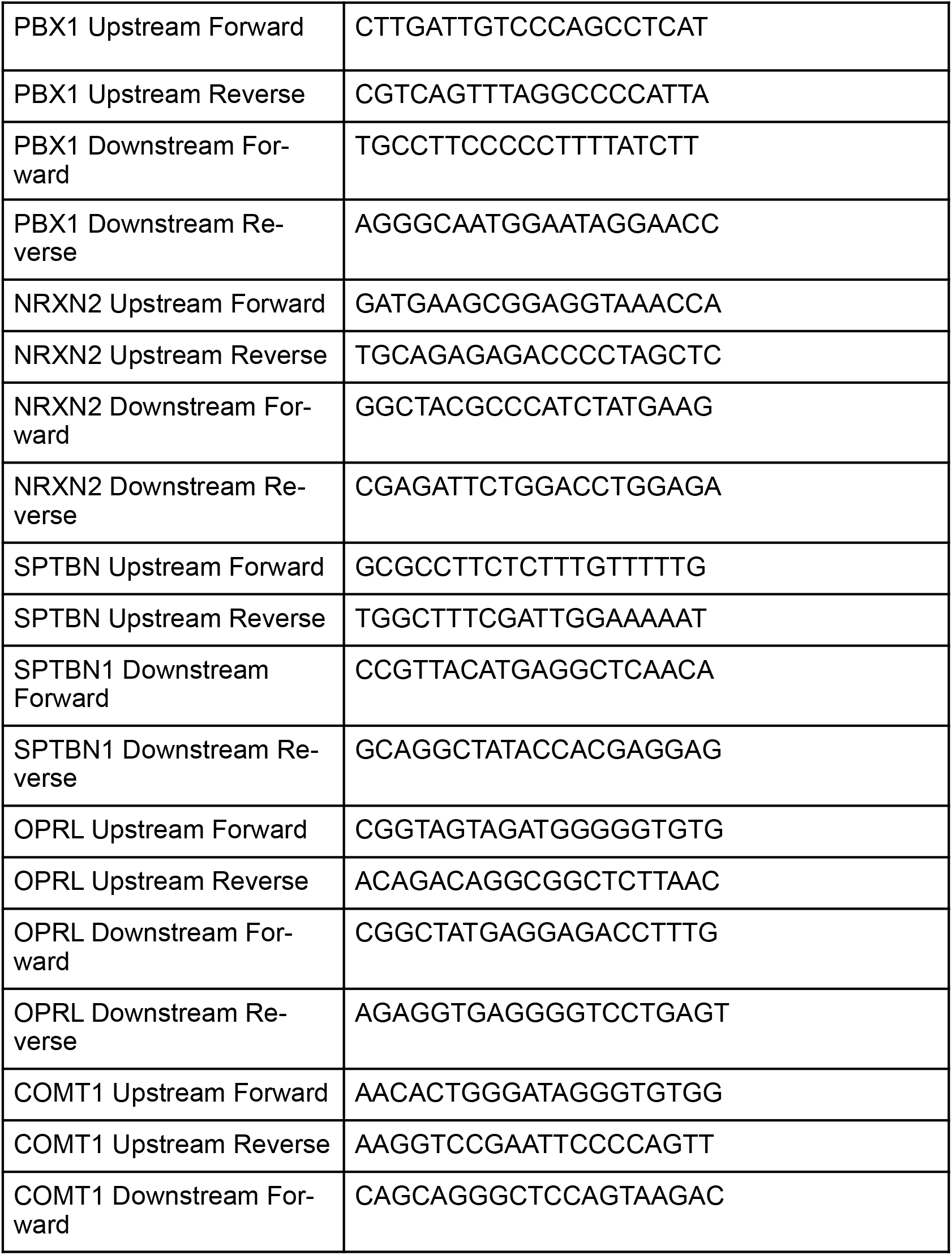
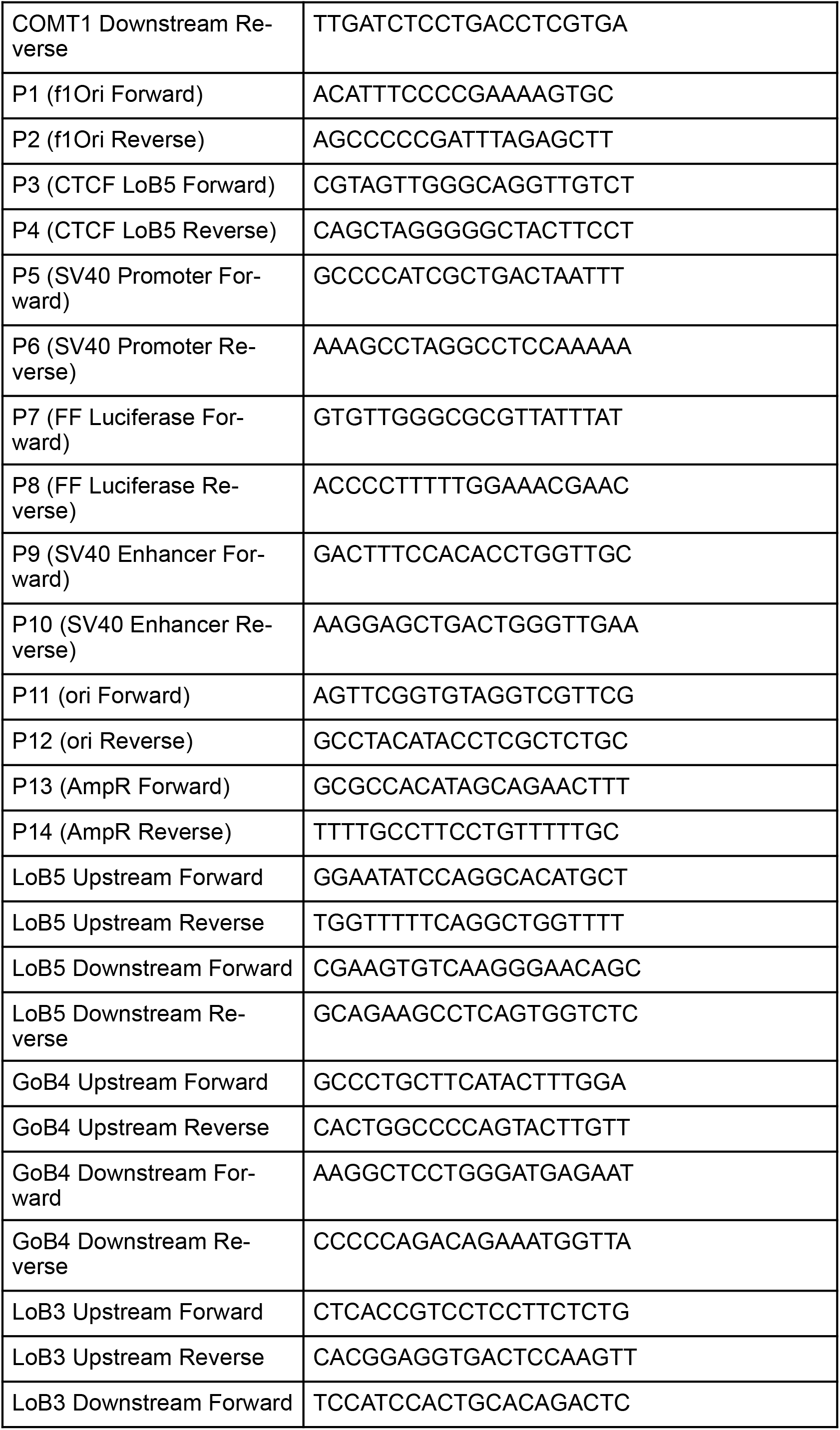
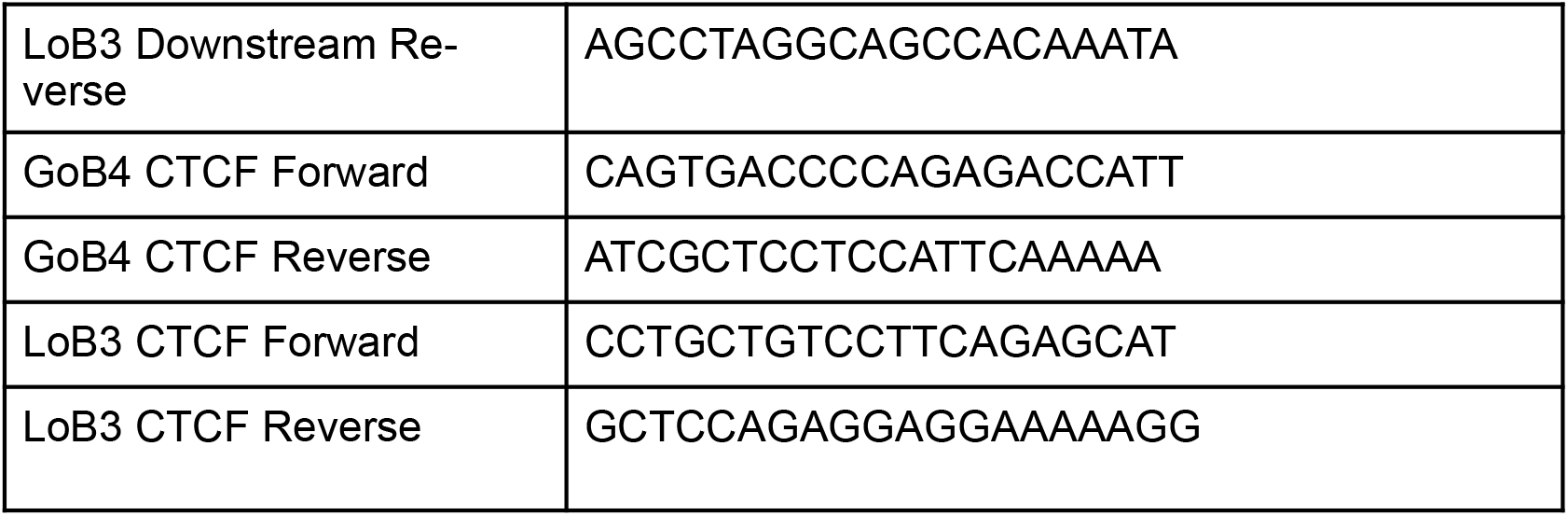

### RNA sequencing

RNA was extracted from CT and KD HEK293T cells. Poly(A)-tailing of RNA was performed using E. coli Poly(A) Polymerase (NEB# M0276) followed by rRNA depletion using NEBNext rRNA Depletion Kit (NEB#E7405S). The Poly(A) RNA thus obtained was used further for library preparation for sequencing on MinION (Oxford Nanopore Technologies). The library preparation for sequencing was carried out following the manufacturer’s protocol for PCR-cDNA Sequencing Kit (SQK-PCS109). Base calling and quality filtration were done in realtime using Guppy in MinKNOW.

### RNA-seq Analysis

The quality-thresholded reads output by Guppy were subjected to trimming of sequencing adapters using porechop2. The fasta format sequences generated by porechop2 were mapped onto the hg38 genome using Hisat2 with these parameters (*-5 35 -3 35 --sensitive -f --ignore-quals --sensitive; including trimming the ends for 35 bases*). The coordinates of the mapped reads were obtained through conversions of sam to bam (samtools view) followed by sorting (samtools sort) and converting to bed (bedtools bamtobed). The fragments were segregated into plus and minus strands followed by merging (bedtools merge) the fragments to generate contiguous regions. The downstream analysis pipeline of the RNA-seq data is described as the schematic diagram as a supplementary figure (Figure S5).

### Plotting RNA-seq signals

RNA-signal files were generated for CT and KD using deeptools bamCoverage. computeMatrix option was used to generate matrix followed by plotting the average type summary plot (plotProfile) or heatmap (plotHeatmap) functions in deeptools.

## Acknowledgments

The authors duly acknowledge the NCCS Pune for HEK293T cells and Mr Sudeep N Banerjee (ISTF, IITGN) for help with computational resources.

US received support from DST-ICPS (T-357) and DBT (BT/PR15883/BRB/10/1480/2016). The studentship of DP and MP was supported by UGC-NET JRF. SD was supported by MHRD, Government of India.

## REFERENCES

Agarwal P, Enroth S, Teichmann M, Jernberg Wiklund H, Smit A, Westermark B, Singh U. 2016. Growth signals employ CGGBP1 to suppress transcription of Alu-SINEs. Cell Cycle 15: 1558–1571.

Arzate-Mejía RG, Recillas-Targa F, Corces VG. 2018. Developing in 3D: the role of CTCF in cell differentiation. Development 145. http://dx.doi.org/10.1242/dev.137729.

Barkess G, West AG. 2012. Chromatin insulator elements: establishing barriers to set heterochromatin boundaries. Epigenomics 4: 67–80.

Becker JS, Nicetto D, Zaret KS. 2016. H3K9me3-Dependent Heterochromatin: Barrier to Cell Fate Changes. Trends Genet 32: 29.

Benoist C, Chambon P. 1981. In vivo sequence requirements of the SV40 early promotor region. Nature 290: 304–310.

Bulut-Karslioglu A, De La Rosa-Velázquez IA, Ramirez F, Barenboim M, Onishi-Seebacher M, Arand J, Galán C, Winter GE, Engist B, Gerle B, et al. 2014. Suv39h-dependent H3K9me3 marks intact retrotransposons and silences LINE elements in mouse embryonic stem cells. Mol Cell 55: 277–290.

Byrne BJ, Davis MS, Yamaguchi J, Bergsma DJ, Subramanian KN. 1983. Definition of the simian virus 40 early promoter region and demonstration of a host range bias in the enhancement effect of the simian virus 40 72-base-pair repeat. Proc Natl Acad Sci U S A 80: 721–725.

Cardiello JF, Kugel JF, Goodrich JA. 2014. A new twist on cell growth control. Cell Cycle 13:3474–3475.

Cuddapah S, Jothi R, Schones DE, Roh T-Y, Cui K, Zhao K. 2009. Global analysis of the insulator binding protein CTCF in chromatin barrier regions reveals demarcation of active and repressive domains. Genome Res 19: 24–32.

Dekker J, Mirny L. 2016. The 3D Genome as Moderator of Chromosomal Communication. Cell 164: 1110–1121.

Farrell CM, West AG, Felsenfeld G. 2002. Conserved CTCF Insulator Elements Flank the Mouse and Human β-Globin Loci. Molecular and Cellular Biology 22: 3820–3831. http://dx.doi.org/10.1128/mcb.22.11.3820-3831.2002.

Galupa R, Crocker J. 2020. Enhancer–Promoter Communication: Thinking Outside the TAD. Trends in Genetics 36: 459–461. http://dx.doi.org/10.1016/j.tig.2020.04.002.

Ghirlando R, Felsenfeld G. 2016. CTCF: making the right connections. Genes Dev 30: 881–891.

Gidoni D, Kadonaga JT, Barrera-Saldaña H, Takahashi K, Chambon P, Tjian R. 1985. Bidirectional SV40 transcription mediated by tandem Sp1 binding interactions. Science 230: 511–517.

Giorgetti L, Lajoie BR, Carter AC, Attia M, Zhan Y, Xu J, Chen CJ, Kaplan N, Chang HY, Heard E, et al. 2016. Structural organization of the inactive X chromosome in the mouse. Nature 535: 575–579.

Gruss C, Wetzel E, Baack M, Mock U, Knippers R. 1988. High-affinity SV40 T-antigen binding sites in the human genome. Virology 167: 349–360.

Hahn M, Dambacher S, Dulev S, Kuznetsova AY, Eck S, Wörz S, Sadic D, Schulte M, Mallm J-P, Maiser A, et al. 2013. Suv4-20h2 mediates chromatin compaction and is important for cohesin recruitment to heterochromatin. Genes Dev 27: 859–872.

Han L, Lee D-H, Szabó PE. 2008. CTCF Is the Master Organizer of Domain-Wide Allele-Specific Chromatin at the H19/Igf2 Imprinted Region. Molecular and Cellular Biology 28: 1124–1135. http://dx.doi.org/10.1128/mcb.01361-07.

Hein MY, Hubner NC, Poser I, Cox J, Nagaraj N, Toyoda Y, Gak IA, Weisswange I, Mansfeld J, Buchholz F, et al. 2015. A human interactome in three quantitative dimensions organized by stoichiometries and abundances. Cell 163: 712–723.

Hertz GZ, Mertz JE. 1988. The enhancer elements and GGGCGG boxes of SV40 provide similar functions in bidirectionally promoting transcription. Virology 163: 579–590.

Holwerda SJB, de Laat W. 2013. CTCF: the protein, the binding partners, the binding sites and their chromatin loops. Philos Trans R Soc Lond B Biol Sci 368: 20120369.

Hou C, Dale R, Dean A. 2010. Cell type specificity of chromatin organization mediated by CTCF and cohesin. Proceedings of the National Academy of Sciences 107: 3651–3656. http://dx.doi.org/10.1073/pnas.0912087107.

Ichiyanagi K. 2014. Regulating Pol III transcription to change Pol II transcriptome. Cell Cycle 13: 3625–3626.

Iglesias N, Moazed D. 2017. Heterochromatin: Silencing repetitive DNA. https://elife-sciences.org/articles/29503 (Accessed November 8, 2020).

Kadesch T, Berg P. 1986. Effects of the position of the simian virus 40 enhancer on expression of multiple transcription units in a single plasmid. Mol Cell Biol 6: 2593–2601.

Kelly JJ, Wildeman AG. 1991. Role of the SV40 enhancer in the early to late shift in viral transcription. Nucleic Acids Research 19: 6799–6804. http://dx.doi.org/10.1093/nar/19.24.6799.

Kim YJ, Cecchini KR, Kim TH. 2011. Conserved, developmentally regulated mechanism couples chromosomal looping and heterochromatin barrier activity at the homeobox gene A locus. Proc Natl Acad Sci U S A 108: 7391–7396.

Klenova EM, Nicolas RH, Paterson HF, Carne AF, Heath CM, Goodwin GH, Neiman PE, Lobanenkov VV. 1993. CTCF, a conserved nuclear factor required for optimal transcriptional activity of the chicken c-myc gene, is an 11-Zn-finger protein differentially expressed in multiple forms. Mol Cell Biol 13: 7612–7624.

Kurukuti S, Tiwari VK, Tavoosidana G, Pugacheva E, Murrell A, Zhao Z, Lobanenkov V, Reik W, Ohlsson R. 2006. CTCF binding at the H19 imprinting control region mediates maternally inherited higher-order chromatin conformation to restrict enhancer access to Igf2. Proc Natl Acad Sci U S A 103: 10684–10689.

Lobanenkov VV, Nicolas RH, Adler VV, Paterson H, Klenova EM, Polotskaja AV, Goodwin GH. 1990. A novel sequence-specific DNA binding protein which interacts with three regularly spaced direct repeats of the CCCTC-motif in the 5’-flanking sequence of the chicken c-myc gene. Oncogene 5: 1743–1753.

Lu Y, Shan G, Xue J, Chen C, Zhang C. 2016. Defining the multivalent functions of CTCF from chromatin state and three-dimensional chromatin interactions. Nucleic Acids Res 44: 6200–6212.

Narendra V, Bulajić M, Dekker J, Mazzoni EO, Reinberg D. 2016. CTCF-mediated topological boundaries during development foster appropriate gene regulation. Genes Dev 30: 2657–2662.

Nichols MH, Corces VG. 2015. A CTCF Code for 3D Genome Architecture. Cell 162: 703–705.

Ninova M, Fejes Tóth K, Aravin AA. 2019. The control of gene expression and cell identity by H3K9 trimethylation. Development 146. http://dx.doi.org/10.1242/dev.181180.

Nishana M, Ha C, Rodriguez-Hernaez J, Ranjbaran A, Chio E, Nora EP, Badri SB, Kloetgen A, Bruneau BG, Tsirigos A, et al. 2020. Defining the relative and combined contribution of CTCF and CTCFL to genomic regulation. Genome Biol 21: 1–34.

Ong C-T, Corces VG. 2014. CTCF: an architectural protein bridging genome topology and function. Nature Reviews Genetics 15: 234–246. http://dx.doi.org/10.1038/nrg3663.

Patel D, Patel M, Datta S, Singh U. 2019. CGGBP1 regulates CTCF occupancy at repeats. Epigenetics & Chromatin 12.http://dx.doi.org/10.1186/s13072-019-0305-6.

Patel D, Patel M, Westermark B, Singh U. 2018. Dynamic bimodal changes in CpG and non-CpG methylation genome-wide upon CGGBP1 loss-of-function. BMC Research Notes 11. http://dx.doi.org/10.1186/s13104-018-3516-1.

Patel M, Patel D, Datta S, Singh U. 2020. CGGBP1-regulated cytosine methylation at CTCF-binding motifs resists stochasticity. BMC Genet 21: 84.

Phillips JE, Corces VG. 2009. CTCF: Master Weaver of the Genome. Cell 137: 1194–1211.http://dx.doi.org/10.1016/j.cell.2009.06.001.

Ren G, Zhao K. 2019. CTCF and cellular heterogeneity. Cell & Bioscience 9. http://dx.-doi.org/10.1186/s13578-019-0347-2.

Sassone-Corsi P, Dougherty JP, Wasylyk B, Chambon P. 1984. Stimulation of in Vitro Transcription from Heterologous Promoters by the SV40 Enhancer. Transfer ∥ Expression of Eukaryotic Genes 7–21. http://dx.doi.org/10.1016/b978-0-12-284650-2.50008-0.

Schmidt D, Schwalie PC, Wilson MD, Ballester B, Gonçalves A, Kutter C, Brown GD, Marshall A, Flicek P, Odom DT. 2012. Waves of retrotransposon expansion remodel genome organization and CTCF binding in multiple mammalian lineages. Cell 148: 335–348.

Shaw PE, Bohmann D, Sergeant A. 1985. The SV40 enhancer influences viral late transcription in vitro and in vivo but not on replicating templates. The EMBO Journal 4: 3247–3252. http://dx.doi.org/10.1002/j.1460-2075.1985.tb04073.x.

Singh U, Roswall P, Uhrbom L, Westermark B. 2011. CGGBP1 regulates cell cycle in cancer cells. BMC Mol Biol 12: 28.

Singh U, Westermark B. 2015. CGGBP1—an indispensable protein with ubiquitous cytoprotective functions. Upsala Journal of Medical Sciences 120: 219–232. http://dx.doi.org/10.3109/03009734.2015.1086451.

Ulaner GA, Yang Y, Hu J-F, Li T, Vu TH, Hoffman AR. 2003. CTCF Binding at the Insulin-Like Growth Factor-II (IGF2)/H19 Imprinting Control Region Is Insufficient to Regulate IGF2/H19 Expression in Human Tissues. Endocrinology 144: 4420–4426. http://dx.doi.org/10.1210/en.2003-0681.

Valadez-Graham V. 2004. CTCF-dependent enhancer blockers at the upstream region of the chicken -globin gene domain. Nucleic Acids Research 32: 1354–1362. http://dx.doi.org/10.1093/nar/gkh301.

Van Bortle K, Ramos E, Takenaka N, Yang J, Wahi JE, Corces VG. 2012. Drosophila CTCF tandemly aligns with other insulator proteins at the borders of H3K27me3 domains. Genome Research 22: 2176–2187. http://dx.doi.org/10.1101/gr.136788.111.

van Kruijsbergen I, Hontelez S, Elurbe DM, van Heeringen SJ, Huynen MA, Veenstra GJC. 2017. Heterochromatic histone modifications at transposons in Xenopus tropicalis embryos. Dev Biol 426: 460.

Wasylyk B, Wasylyk C, Augereau P, Chambon P. 1983. The SV40 72 bp repeat preferentially potentiates transcription starting from proximal natural or substitute promoter elements. Cell 32: 503–514.

Weintraub AS, Li CH, Zamudio AV, Sigova AA, Hannett NM, Day DS, Abraham BJ, Cohen MA, Nabet B, Buckley DL, et al. 2017. YY1 Is a Structural Regulator of Enhancer-Promoter Loops. Cell 171: 1573.

Yusufzai TM, Tagami H, Nakatani Y, Felsenfeld G. 2004. CTCF Tethers an Insulator to Sub-nuclear Sites, Suggesting Shared Insulator Mechanisms across Species. Molecular Cell 13: 291–298. http://dx.doi.org/10.1016/s1097-2765(04)00029-2.

