## Supplementary material for "CGGBP1-dependent CTCF-binding sites restrict ectopic transcription": Fig S1

### Figure S1

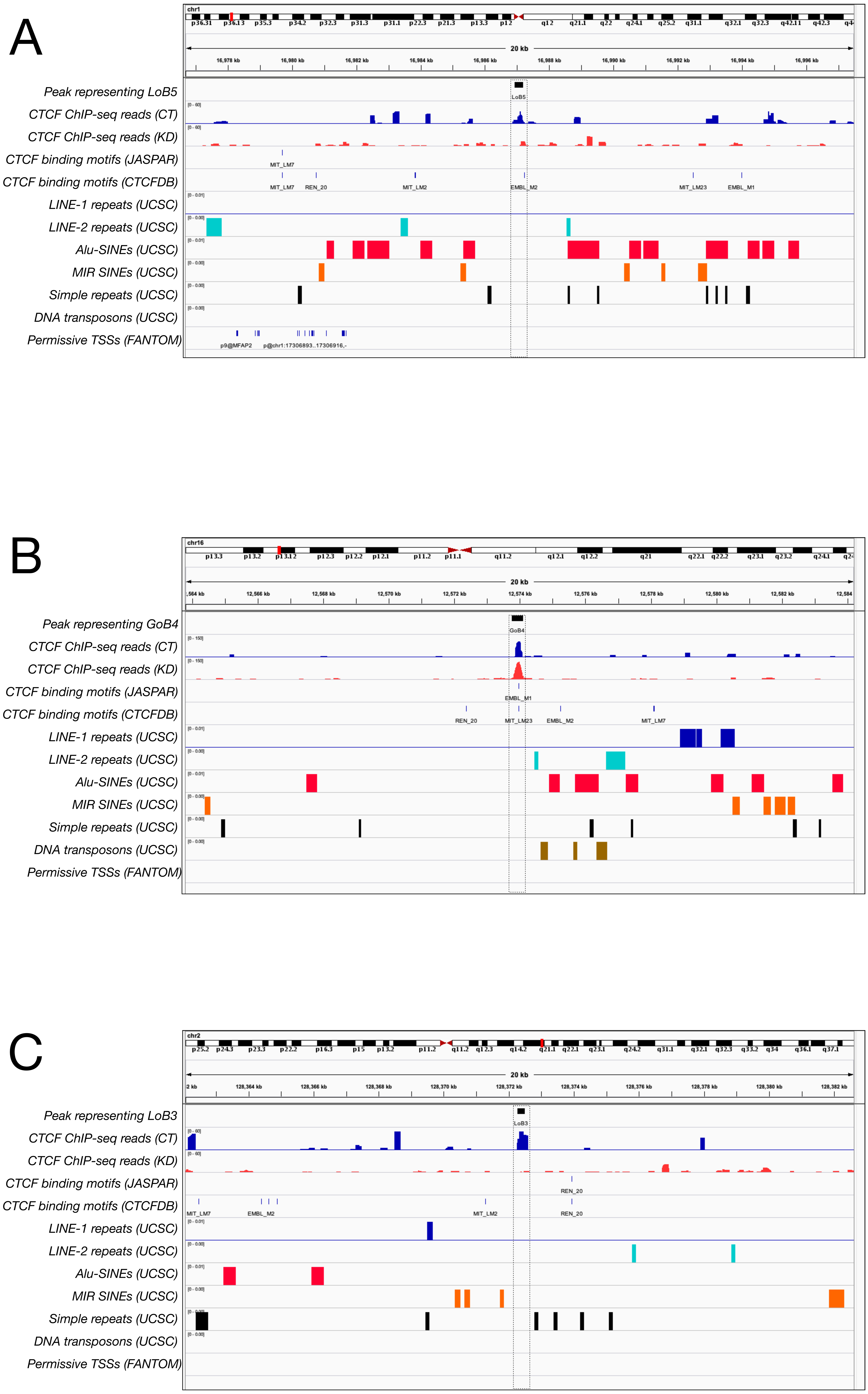


Fig S1: Characterisation of CGGBP1-regulated CTCF binding sites for repeats and CTCF-motif. **A-C:** Genome browser views showing the distribution of CTCF ChIP-seq reads, predicted CTCF binding motifs by MEME-suite FIMO and CTCFBSDB and permissive TSSs. Repeat tracks show the distribution of the LINE-1, LINE-2, Alu, MIR-SINEs, Simple repeats and DNA-repeat elements. **A:** The genome browser view on chromosome 1 (~Chr1:16976 kb - 16996 kb) shows loss of CTCF ChIP-signal at LoB5 upon CGGBP1 depletion. Alu-SINEs and MIR-SINEs are clustered in immediate flanks of the LoB5 CTCF binding sites. The nearest permissive TSS is approximately 8 kb upstream from the LoB5 site. **B:** The genome browser view on chromosome 16 (~Chr16:12564 kb - 12584 kb) represents GoB4 CTCF binding site. CTCF ChIP-signal is potentiated in KD at the GoB4 site. Predicted CTCF motifs (EMBL_M1 and MIT_LM23) are present in the GoB4 CTCF binding site. Interspersed repeats (LINE-1, LINE-2, Alu and MIR-SINEs) and DNA-elements are clustered downstream of the GoB4 CTCF binding site. **C:** The genome browser view on chromosome 2 (~Chr2:128362 kb - 128382 kb) represents LoB3 CTCF binding site. CTCF ChIP-signal is reduced at LoB3 CTCF binding site. SINEs (Alu and MIR-SINEs) and DNA repeats are enriched in the upstream and downstream of the LoB3 CTCF binding site respectively.
