## Supplementary material for "CGGBP1-dependent CTCF-binding sites restrict ectopic transcription": Fig S2

Figure S2

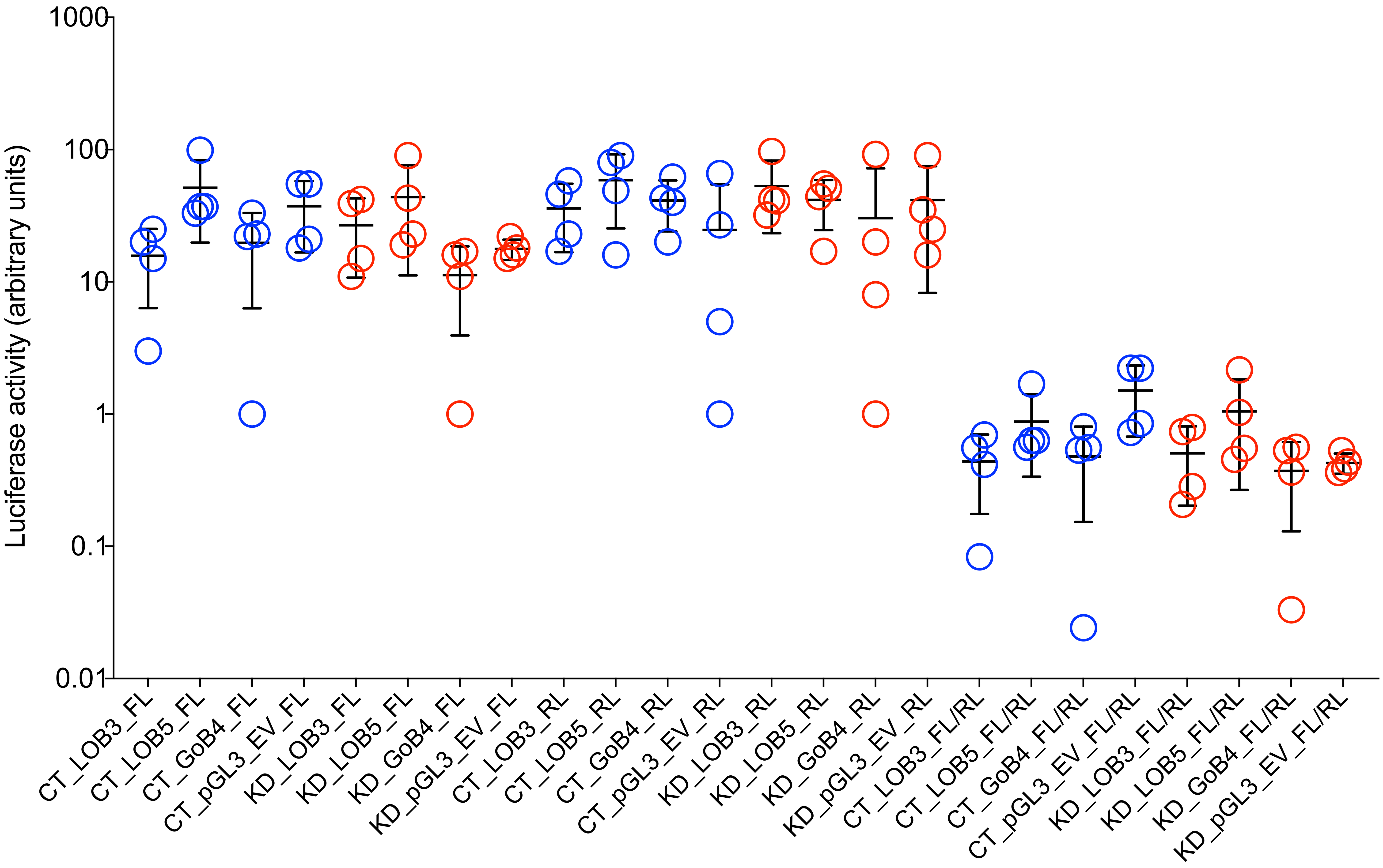


Fig S2: Luciferase reporter gene assays. For the three episomal constructs LoB3, LoB5 and GoB4, the Luciferase reporter gene activity was determined to measure the *cis*-regulatory effect of the SV40 promoter (juxtaposed with LoB3, LoB5 or GoB4 respectively). The luminescence values for each sample (X-axis) is reported on the Y-axis (log 10 scale). Values are from separate experiments and account for multiple systemic experimental variations. The values shown are mean and standard deviations. The suffixes FL, RL and FL/RL stand for luminescence values of Firefly Luciferase, Renilla Luciferase and the ratio of Firefly Luciferase to Renilla Luciferase respectively. Renilla Luciferase values serve as an internal control.
