## Supplementary material for "CGGBP1-dependent CTCF-binding sites restrict ectopic transcription": Fig S3

### Figure S3

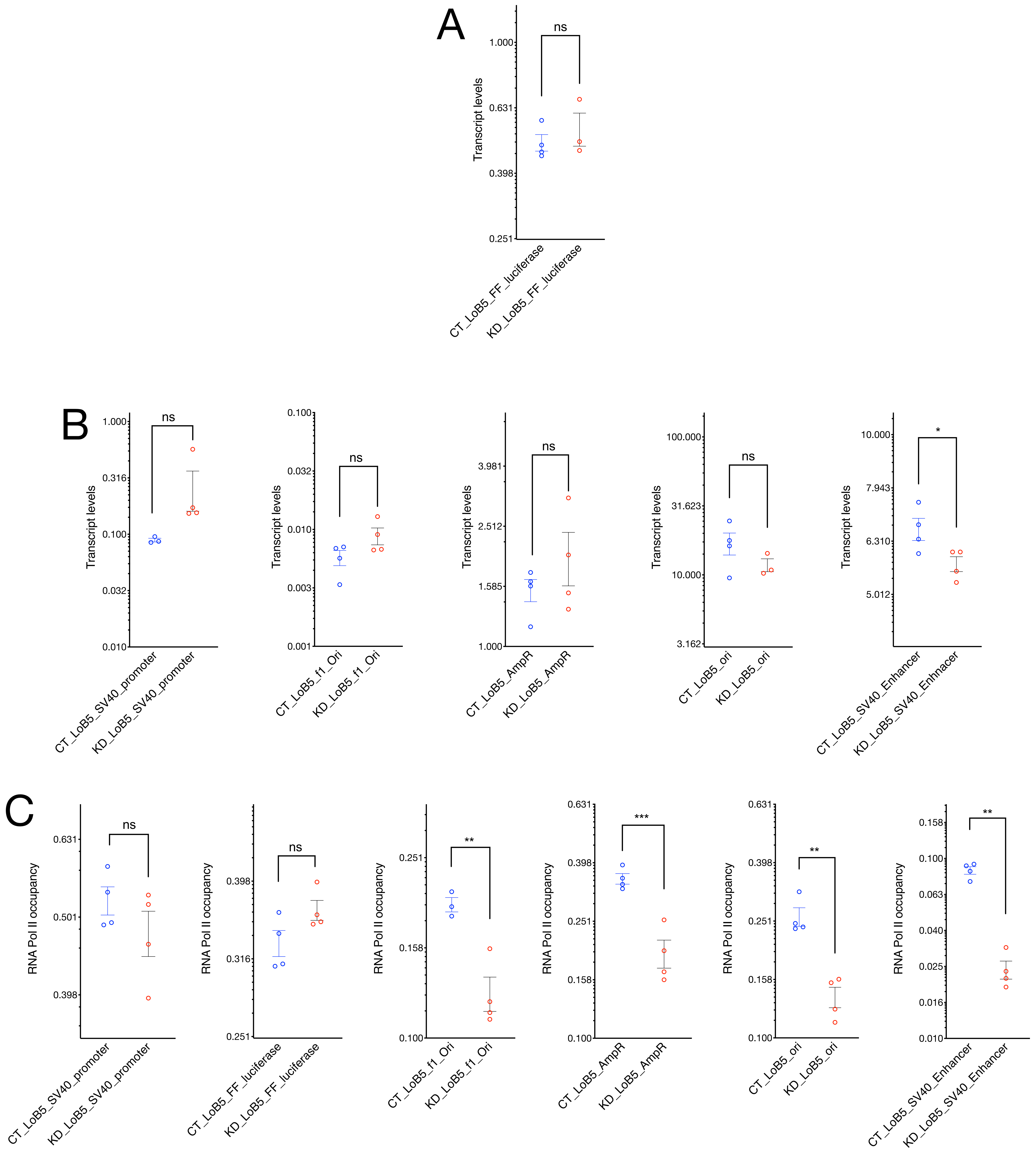


Fig S3: Regulation of the LoB5-SV40 promoter activity by CGGBP1. **A:** pGL3-LoB5 construct was transfected in HEK293T cells with normal and depleted levels of CGGBP1. Transcript levels were compared at different episomal regions between CT and KD. SV40 Promoter and FF luciferase show a non-significant increase in transcript levels upon removal of CGGBP1. **B:** Similarly, SV40 promoter, immediate upstream located f1Ori and AmpR regions also show a non-significant increase in transcript levels in the absence of CGGBP1. However, the SV40 enhancer exhibits a strong decline in transcriptional activity, while Ori remains immune to any significant transcriptional changes upon CGGBP1 depletion. **C:** RNA Polymerase II occupancy was compared across the episomal landscape between CT and KD. In agreement with transcript levels, the SV40 promoter and FF luciferase do not show significant changes in RNA polymerase II levels. However, f1Ori and AmpR, the immediate upstream regions of the episomal LoB5 site, show a significant decrease in RNA Polymerase II levels in KD. Similarly, RNA Polymerase II levels at SV40 enhancer and Ori portray a significant reduction in KD.
