## Supplementary material for "CGGBP1-dependent CTCF-binding sites restrict ectopic transcription": Fig S4

### Figure S4

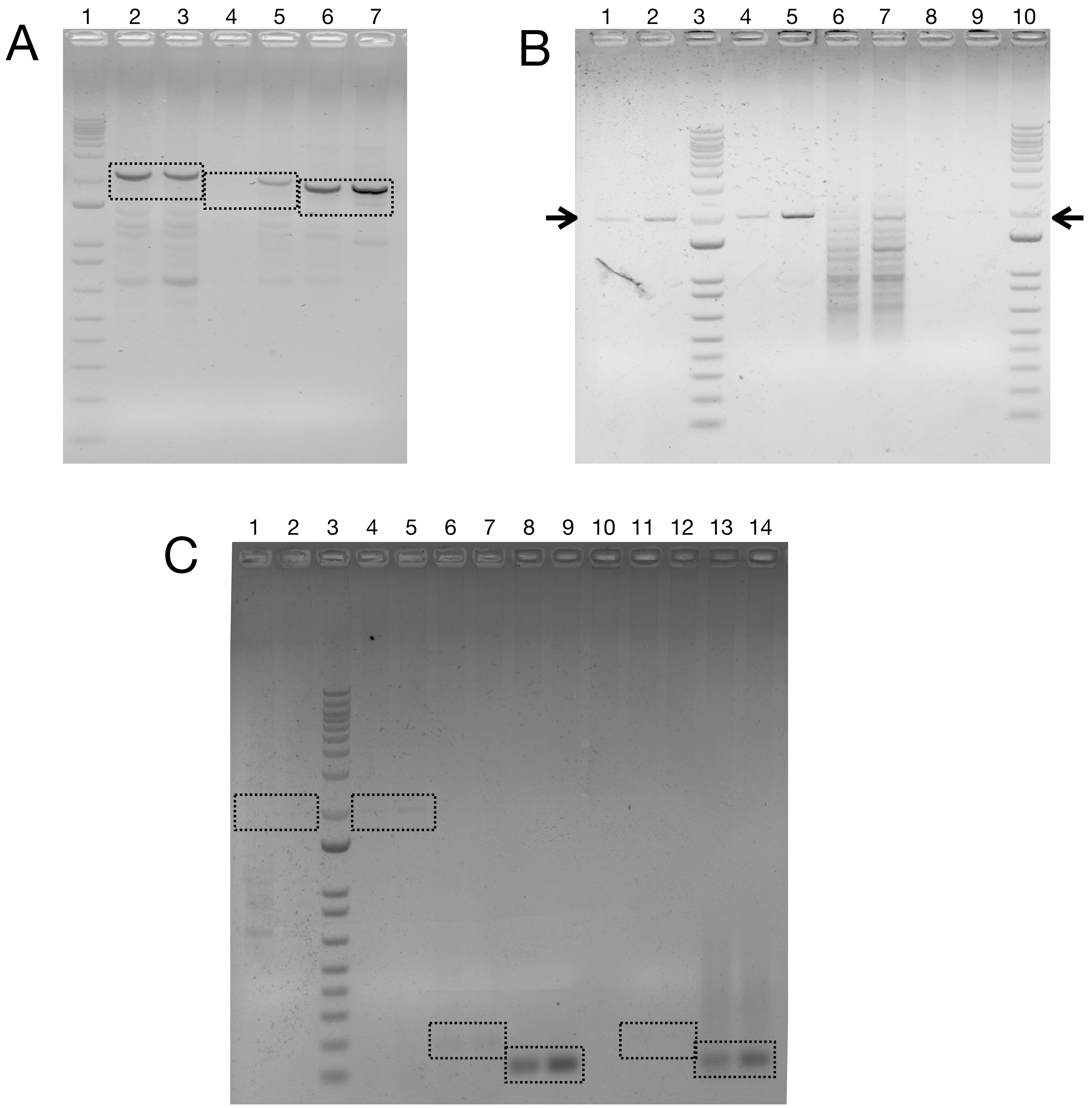


Fig S4: CGGBP1 regulates atypical promoter activity of the SV40 promoter in the pGL3-Control vector system. **A:** Randomly primed cDNA from LoB3, LoB5 and GoB4 pGL3-control construct transfected CT and KD cells were used to detect transcripts spanning CGGBP1-regulated CTCF binding sites by using primer P1 (f1Ori Forward) and P6 (SV40 Promoter Reverse). The strongest difference in transcript levels was observed in LoB5 followed by GoB4. Lane order is as follows: 1: DNA ladder, 2 and 3: cDNA using random primers and PCR using P1 and P6; CT (2) and KD (3); plasmid pGL-Control-LoB3, 4 and 5: cDNA using random primers and PCR using P1 and P6; CT (4) and KD (5); plasmid pGL-Control-LoB5, 6 and 7: cDNA using random primers and PCR using P1 and P6; CT (6) and KD (7); plasmid pGL-Control-GoB4. The expected molecular weight of PCR products (2.15 kb for LoB3 lanes 2 and 3, 1.97 kb for LoB5 lanes 4 and 5, and 1.86 kb for GoB4, lanes 6 and 7) are indicated by boxes. **B:** RNA was isolated from pGL3-control LoB5 construct transfected cells. To study atypical promoter activity of the LoB5-SV40 promoter strand-specific cDNA were synthesised from DNaseI digested RNA by using promoter P1 (f1Ori Forward) and P6 (SV40 Promoter Reverse) primers. The full-length 1.97 kb PCR product was detected using oligo-dT primed cDNA as well as P1 primed cDNA but not P6 primed cDNA. For oligo-dT as well as P1 primed cDNA, the levels of this transcript were higher in KD than in CT. Using P6 primed cDNA multiple non-specific fragments of smaller molecular weights were generated. As a control, the DNase-digested RNA template did not yield any PCR amplification. Lane order is as follows: 1 and 2: cDNA using oligo-dT, PCR using P1 and P6; CT (1) and KD (2), 3: DNA ladder, 4 and 5: cDNA using P1 and PCR using P1 and P6; CT (4) and KD (5), 6 and 7: cDNA using P6 and PCR using P1 and P6; CT (6) and KD (7), 8 and 9: RNA (pGL3-Control-LoB5 transfected), PCR using P1 and P6; CT (8) and KD (9), 10: DNA ladder. The expected molecular weight of PCR products (1.97 kb) are indicated by an arrow. **C:** RNA was isolated from pGL3-control LoB5 construct transfected and untransfected cells. To study atypical promoter activity of the LoB5-SV40 promoter strand-specific cDNA were synthesised from DNaseI digested RNA by using promoter P1 (f1Ori Forward) and P6 (SV40 Promoter Reverse). As expected, no specific PCR product (1.97 Kb) was detected by using RNA as a template from untransfected CT and KD cells. However, low levels of the specific PCR product were observed by using DNaseI digested RNA from LoB5 pGL3 control construct transfected CT and KD cells. The specific amplification was comparatively higher in KD. Such a weak amplification may be due to the weak reverse transcriptase activity of the Taq DNA Polymerase. Strand Specific transcriptions were compared at f1Ori and SV40 Promoter. Very low transcriptional levels at f1Ori were observed as compared to SV40 promoter by using cDNA from P1 (f1Ori Forward). KD has shown higher transcript levels compared to CT at SV40 promoter and f1Ori. Similarly, no specific PCR products were observed at f1Ori and weak transcriptional activity was observed as SV40 promoter by using cDNA from P6 (SV40 Promoter Reverse). Lane order is as follows: 1 and 2: RNA (no pGL3-Control-LoB5), PCR using P1 and P6; CT (1) and KD (2), 3: DNA ladder, 4 and 5: RNA (pGL3-Control-LoB5 transfected), PCR using P1 and P6; CT (4) and KD (5), 6 and 7: cDNA using P1 and PCR using P1 and P2; CT (6) and KD (7), 8 and 9: cDNA using P1 and PCR using P5 and P6; CT (8) and KD (9), 10: Empty lane, 11 and 12: cDNA using P6 and PCR using P1 and P2; CT (11) and KD (12), 13 and 14: cDNA using P6 and PCR using P5 and P6; CT (13) and KD (14). The expected molecular weight of PCR products (1.97 kb) are indicated by boxes.
