## Supplementary figures and images for "CGGBP1-dependent CTCF-binding sites restrict ectopic transcription"

### Fig S5

#### Figure S5

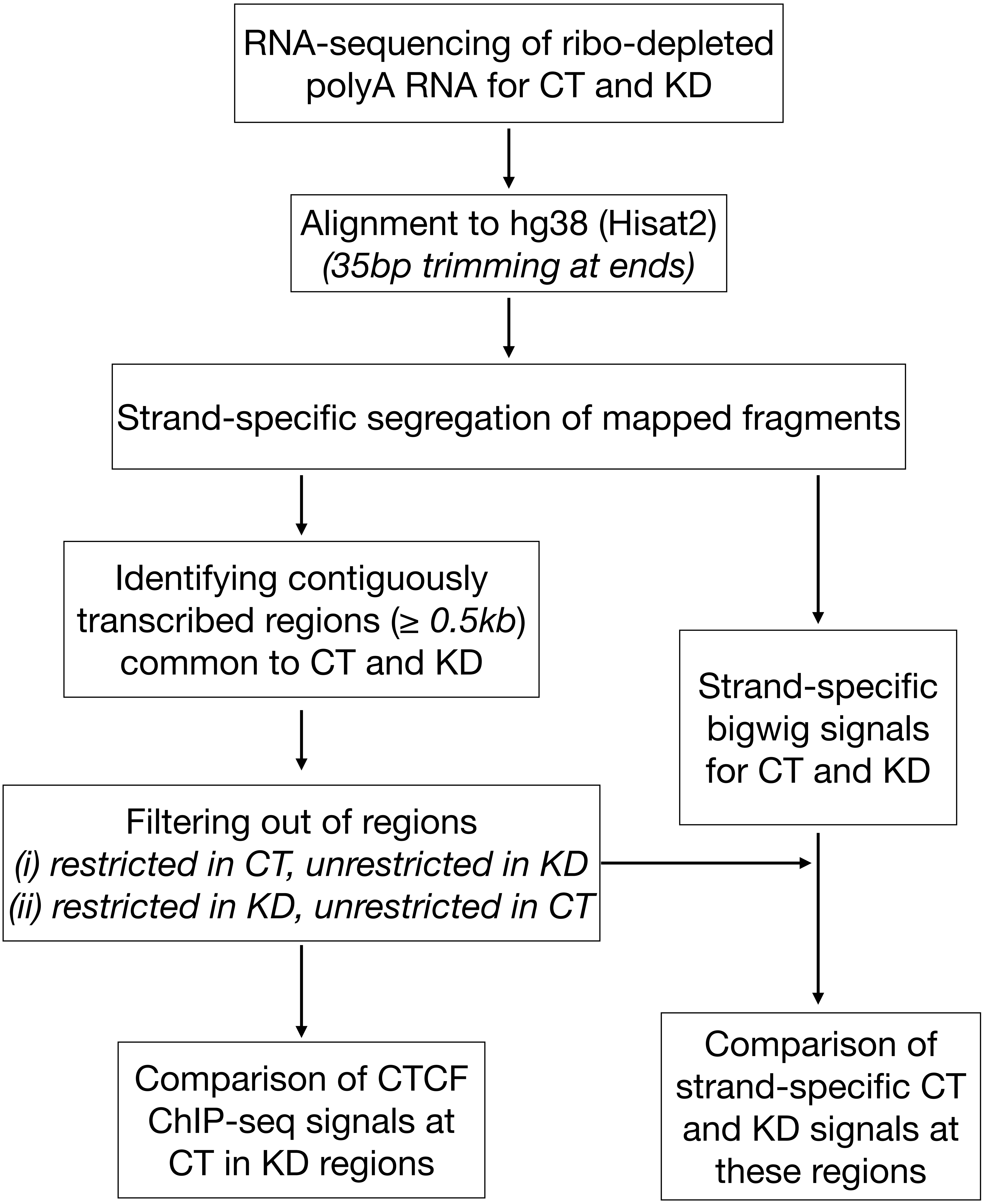


Fig S5: A schematic representation of RNA-seq data analysis pipeline.
